# Cell-type specific deletions of Neuroligin 2 reveal a vital role of synaptic excitation-inhibition balance

**DOI:** 10.1101/2022.09.23.509267

**Authors:** Colleen M. Longley, Xize Xu, Jessica E. Messier, Zhao-Lin Cai, Jung Woo Park, Hongmei Chen, Derek L. Reznik, Monika P. Jadi, Mingshan Xue

## Abstract

Synaptic excitation (E) and inhibition (I) stay relatively proportional to each other over different spatiotemporal scales, orchestrating neuronal activity in the brain. This proportionality, referred to as E-I balance, is thought to be critical for neuronal functions because its disruption was observed in many neurological disorders. However, the causal evidence demonstrating its significance is scarce. Here we show that deleting Neuroligin-2 (Nlgn2), a postsynaptic adhesion molecule at inhibitory synapses, from mouse glutamatergic or GABAergic neurons reduces inhibition cell-autonomously without affecting excitation, thereby disrupting E-I balance and causing lethality. In contrast, deleting Nlgn2 constitutively or simultaneously from both glutamatergic and GABAergic neurons results in viable mice. A neural network model shows that reducing inhibition in either neuronal type is detrimental to network activity, but in both types partially re-establishes E-I balance and activity. Together, our results provide evidence for an essential role of E-I balance in brain functions and organism survival.

## Introduction

Cortical networks rely on a precise balance between excitation (E) and inhibition (I) to ensure efficient information processing (Haider and Mccormick, 2009). In a single neuron, the synaptic weights from E and I are co-tuned such that a cell that receives more excitation correspondingly receives more inhibition as well through homeostatic plasticity (Xue, Atallah and Scanziani, 2014). Meanwhile, in the entire network E and I are dynamically balanced such that the mean inhibition and mean excitation received by neurons is balanced across different patterns of activity without directly altering synaptic weights (Ahmadian and Miller, 2019). Additionally, many cortical regions, including visual cortex, operate as inhibition-stabilized networks, meaning that as external excitatory input to the network increases due to sensory inputs, the feedback inhibition increases as well, allowing the network to stabilize (Adesnik, 2017).

The importance of E-I balance is demonstrated by its role in the pathogenesis of many neurological disorders, including Autism Spectrum Disorder (ASD) (Rubenstein and Merzenich, 2003). However, the causal evidence linking E-I balance and ASD is scarce. It has been shown that acutely disrupting E-I balance in a wild-type mouse using optogenetics leads to the development of ASD symptoms ((Yizhar *et al*., 2011) while acutely restoring E-I balance using optogenetics in a mouse model of ASD is able to alleviate ASD symptoms (Selimbeyoglu *et al*., 2017). However, these are acute manipulations and they do not address the role of E-I balance during development. It has been demonstrated that increasing or decreasing sensory input during the critical window leads to the development of more noisy networks that are more prone to the generation of seizures (Zhang, Bao and Merzenich, 2001; Chang and Merzenich, 2003), suggesting that E-I balance during development might play an important role in ASD pathogenesis. Additionally, recent work has suggested that the E-I imbalance observed in ASD mouse models may be compensatory, rather than causative, as it serves to restore normal neural activity rather than alter it (Antoine *et al*., 2019).

There is mounting evidence that there is not only a disruption of E-I balance underlying ASD pathogenesis, but more specifically, that there is a decrease in the level of inhibition driving an increase in E-I ratio. The importance of inhibitory synapses in ASD is evidenced by the many mouse models disrupting inhibition that display neuropsychiatric symptoms. For example, mice lacking the Gabbr1 gene, which encodes for a subunit of the GABAB receptor, display spontaneous seizures, hyperactivity, and memory impairment (Schuler et al., 2001). It has also been shown in some mouse models of human disease that many of the disease phenotypes can be recapitulated by deleting the causative gene only in inhibitory cells. For example, it has been shown that deletion of Mecp2, the gene that causes Rett syndrome, specifically in GABAergic cells recapitulates most of the phenotypes seen in a constitutive deletion (Chao et al., 2010). In addition to the role of E-I balance and disrupted inhibition in ASD, it has also been implicated in other neuropsychiatric disorders like epilepsy and schizophrenia. Patients with schizophrenia display alterations in gene expression in GABAergic interneurons of the prefrontal cortex that may contribute to some of the cognitive deficits associated with this disease (Akbarian *et al*., 1995; Hashimoto *et al*., 2003). Additionally, epilepsy is frequently characterized by hyper-excitable cortical networks that result from a breakdown of E-I balance. (Dehghani *et al*., 2016). Thus, targeting E-I balance, and particularly GABAergic interneuron activity, is a promising strategy for treating a diverse array of neurodevelopmental and psychiatric disease.

All inhibition in the cortex comes from the activity of GABAergic interneurons which form hyperpolarizing synaptic contacts primarily with other local neurons (Tremblay, Lee and Rudy, 2016). While most of our understanding of inhibition focuses on the interaction between interneurons and excitatory neurons there are also inhibitory synapses that exist between different interneurons (Staiger, Freund and Zilles, 1997; Staiger *et al*., 2004). While the classification of interneuron types is complex (Ascoli *et al*., 2008) they can be broadly sub-classified into three major sub-types based on gene expression: Parvalbumin (Pv) expressing, Somatostatin (Sst) expressing, and 5HT3aR expressing. The 5HT3aR population can be further subclassified as vasoactive-intestinal peptide (Vip) positive and Vip negative (Rudy *et al*., 2011). Sst and Pv positive interneurons provide most of the inhibition onto excitatory pyramidal cells while VIP interneurons primarily provide inhibition onto Sst interneurons, functioning to disinhibit pyramidal cells (Pfeffer *et al*., 2013; Jiang *et al*., 2015). Despite recent work on the functional impact of these disinhibitory connections (Xu *et al*., 2019), the differential contributions of these two classes of inhibitory synapses to E-I balance is not known. One way to differentially manipulate these two classes of inhibitory synapses while also disrupting E-I balance during development is to selectively delete key components of the inhibitory post-synapse in either excitatory or inhibitory neurons.

Neuroligin 2 (Nlgn2) is a post-synaptic cell adhesion molecule that is exclusively expressed at inhibitory synapses (Varoqueaux, Jamain and Brose, 2004). Loss of Nlgn2 disrupts inhibitory synaptic activity but has no effect on excitatory synaptic activity (Chubykin *et al*., 2007). It is thought to participate in the recruitment of the GABA_A_ receptor to the post-synaptic site by interactions with pre-synaptic neurexin, gephyrin, and others (Poulopoulos *et al*., 2009). While much is known about the cellular and behavioral effects of constitutive Nlgn2 knock-out (KO) (Blundell *et al*., 2009; Gibson, Huber and Südhof, 2009; Babaev *et al*., 2016; Seok *et al*., 2018; Cao *et al*., 2020) relatively little is known about the cell-autonomous effects of conditional Nlgn2 in different cell types (Liang *et al*., 2015; Zhang *et al*., 2015; Zhang and Südhof, 2016).

To this end, we investigated the effect of Nlgn2 at inhibitory synapses onto interneurons by performing cell-type specific deletion of Nlgn2 specifically in the different subtypes of GABAergic interneurons. Using interneuron specific Cre lines had no effect on inhibition onto interneurons but targeted viral delivery of iCre into the three classes of interneurons was able to significantly reduce inhibition. Further investigation into the interneuron specific Cre lines revealed that they were unable to achieve complete deletion of Nlgn2 mRNA before the third post-natal week while viral delivery of iCre achieved a much earlier deletion. This suggested that the timing of Nlgn2 deletion is critical. We further verified this by testing the effect of adult deletion of Nlgn2 in interneurons and pyramidal cells and confirmed that adult deletion did not affect inhibition. Knowing that Nlgn2 knock-out can reduce inhibition onto both excitatory and inhibitory neurons we tested the effect on E-I balance by conditionally deleting Nlgn2 in all excitatory or all inhibitory interneurons. Much to our surprise, deletion of Nlgn2 in excitatory neurons is lethal but simultaneously deleting Nlgn2 in inhibitory neurons as well restores viability. We hypothesized that this lethality is due to severe alteration in E-I balance. We investigated this hypothesis using a simple model of a cortical network and showed that decreasing inhibitory inputs onto excitatory cells results in severe instability of the model but simultaneously decreasing inhibitory inputs onto both excitatory and inhibitory neurons restores stability. This suggests that a proper balance between inhibitory synapses onto both excitatory and inhibitory neurons is critical for the survival of the animal, confirming the importance of proper E-I balance. Additionally, our work suggests that using cell-type specific gene knockouts to infer information about gene function may reveal unexpected findings due to the interactions between different cell types.

## Results

### Nlgn2 is expressed in all three major subtypes of cortical interneurons

Constitutive Nlgn2 knock-out mice have previously been shown to have decreased inhibitory synaptic input onto numerous different cell types including brainstem excitatory neurons (Poulopoulos *et al*., 2009), cortical pyramidal cells (Gibson, Huber and Südhof, 2009), and hippocampal pyramidal cells (Horn *et al*., 2017). However, the function of Nlgn2 in cortical interneurons was not known. First, we sought to determine whether Nlgn2 is expressed in cortical interneurons. To do this we performed double fluorescent *in situ* hybridization (dFISH) against *Nlgn2* and either *Somatostatin* (Sst), *Parvalbumin* (Pv), or *Vasoactive intestinal peptide* (Vip) to label the different classes of interneurons. We designed a DIG conjugated RNA probe for *Nlgn2* that binds to exons 4-6 of the mouse *Nlgn2* mRNA. Exons 4-6 are the exons that are flanked by LoxP sites and removed by Cre recombinase in the conditional knock-out *Nlgn2* mouse line, so that region should not be present in Cre expressing cells (Liang *et al*., 2015) (Supp Fig 1 A). First, we confirmed the specificity of our *Nlgn2* probe by comparing the Nlgn2 fluorescent intensity in layer 2/3 neurons from visual cortex of WT and Nlgn2^-/-^ animals. We found an 89.2% reduction in the average cellular Nlgn2 fluorescence intensity in Nlgn2^-/-^ animals indicating that our probe is specific for *Nlgn2* (Supp Fig 1 B,C). For Sst interneurons we found that 77.7% of the cells in the WT animals had a higher Nlgn2 fluorescence intensity than the Nlgn2^-/-^ cells (Supp Fig 1 D,E). Similarly, for Vip interneurons we found that 86.9% of the cells in the WT animals were higher than the Nlgn2^-/-^ cells (Supp Fig 1 F,G), and for Pv interneurons we found that 93.7% of the cells in the WT animals were higher than the Nlgn2^-/-^ cells (Supp Fig 1 H,I). Together, this data confirms *Nlgn2* is expressed in all three major subtypes of cortical neurons. This led us to question whether Nlgn2 is also playing a similar role in regulating their inhibition.

### Conditional *Nlgn2* knockout via interneuron specific Cre lines has no effect on inhibitory synaptic inputs

To determine the effect of loss of Nlgn2 on cortical interneurons we first conditionally knocked out Nlgn2 in Sst interneurons by breeding *Nlgn2 floxed* mice with *Sst-ires-cre* mice. We obtained acute coronal brain slices from 3-5 week old *Nlgn2^f/f;^Sst^Cre/+^* mice and their age- and sex-matched *Nlgn2^+/+^;Sst^Cre/+^* littermates injected in visual cortex at P0-P1 with AAV9-DIO-TdTomato to label the Cre+ cells. We performed whole-cell voltage clamp to record spontaneous excitatory postsynaptic currents (sEPSCs) and inhibitory postsynaptic currents (sIPSCs) from TdTomato+ Sst neurons in layer 5. Surprisingly, we found that the frequencies and amplitudes of sEPSCs and sIPSCs in Nlgn2^f/f^;Sst^Cre/+^ mice were not significantly different from the controls (Fig 1 A,B). For Vip interneurons we obtained acute coronal brain slices from 3-5 week old *Nlgn2^f/f^*;*Vip^Cre/+^;Rosa26^Ai14/+^*and *Nlgn2^+/+^;Vip^Cre/^;Rosa26^Ai14/+^* mice and recorded from TdTomato positive neurons in layer 2/3. We only found a small decrease in sIPSC amplitude in *Nlgn2^f/f^*;*Vip^Cre/+^;Rosa26^Ai14/+^*mice as compared to the controls (Fig 1 C,D). For Pv interneurons we obtained acute coronal brain slices from 10-14 week old *Nlgn2^f/f^*;*Pv^Cre/+^;Rosa26^Ai14/+^* and *Nlgn2^+/+^;Pv^Cre/+^;Rosa26^Ai14/+^* mice and recorded from TdTomato+ neurons in layer 5. Again, we found no difference in sEPSCs or sIPSCs in *Nlgn2^+/+^;Pv^Cre/+^;Rosa26^Ai14/+^* mice (Fig 1 E,F).

**Figure 1:**
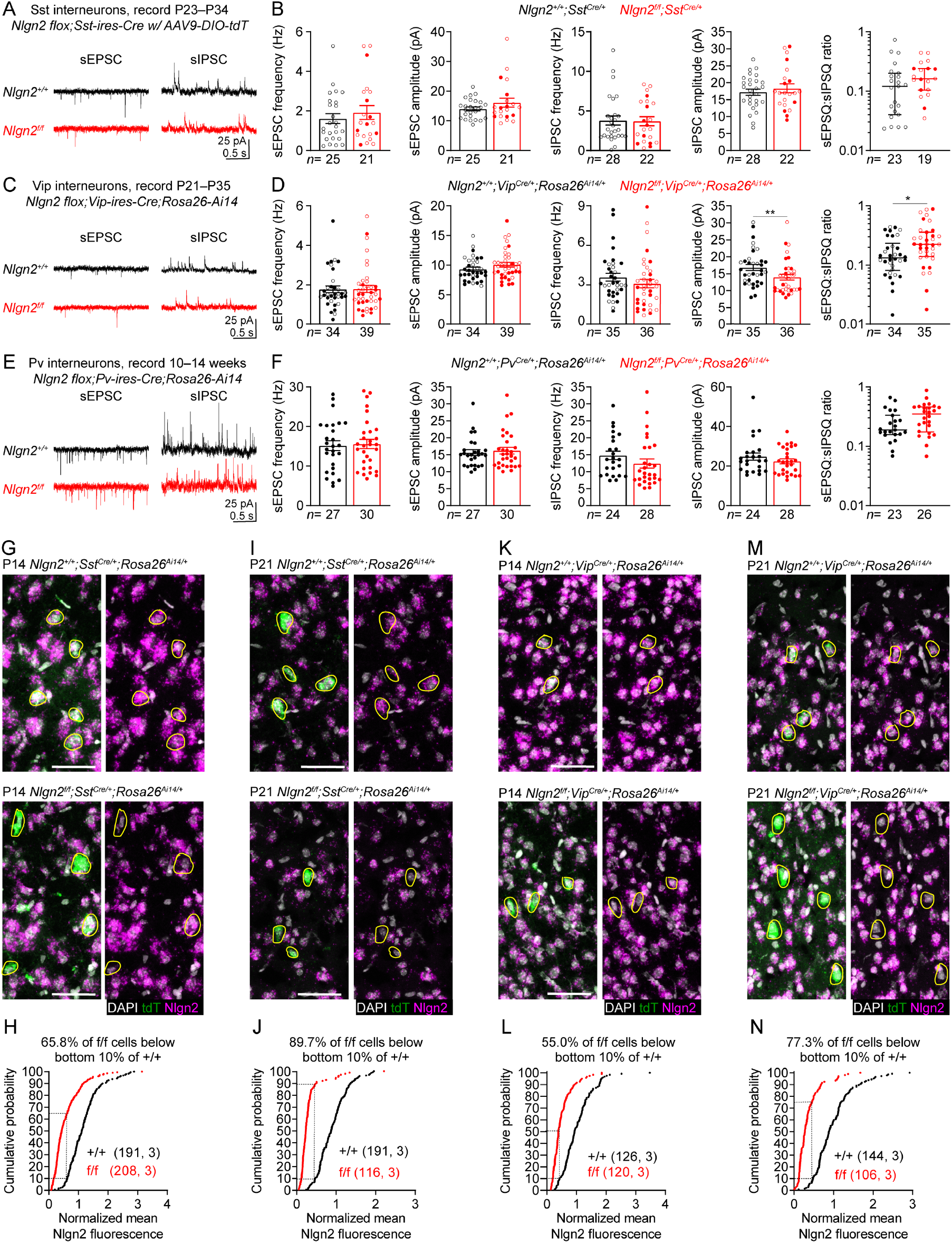
Targeting Nlgn2 with interneuron-specific Cre lines does not affect inhibition (**A**) Representative traces of sEPSCs (left) and sIPSCs (right) of a layer 5 Sst interneuron in the visual cortex from a *Nlgn2^+/+^;Sst^Cre/+^*(top) and *Nlgn2^f/f^*;*Sst^Cre/+^* (bottom) mouse. (**B**) Summary data of the frequency and amplitude of sEPSCs, sIPSCs, and the sEPSQ:sIPSQ ratio between *Nlgn2^+/+^;Sst^Cre/+^*and *Nlgn2^f/f^*;*Sst^Cre/+^* mice. (**C,D**) As in (A,B), but for Vip interneurons in layer 2/3 of *Nlgn2^+/+^;Vip^Cre/^;Rosa26^Ai14/+^*and *Nlgn2^f/f^*;*Vip^Cre/+^;Rosa26^Ai14/+^*mice (**E,F**) As in (A,B), but for Pv interneurons in layer 5 from *Nlgn2^+/+^;Pv^Cre/+^;Rosa26^Ai14/+^*and *Nlgn2^f/f^*;*Pv^Cre/+^;Rosa26^Ai14/+^*mice. (**G**) Representative images of DFISH showing *Nlgn2* (magenta), *tdTomato* (green), and DAPI (white) in the visual cortex of P14 *Nlgn2^+/+^;Sst^Cre/+^;Rosa26^Ai14/+^*(top) and *Nlgn2^f/f^*;*Sst^Cre/+^;Rosa26^Ai14/+^*(bottom) mice. Yellow outlines represent tdTomato positive cells. (**H**) Cumulative frequencies of tdTomato positive cells as a function of their normalized *Nlgn2* levels. Dashed lines indicate that the Nlgn2 levels of 65.8% of tdTomato positive cells from the P14 *Nlgn2^f/f^*;*Sst^Cre/+^;Rosa26^Ai14/+^*are within the bottom 10% of tdTomato positive cells from *Nlgn2^+/+^;Sst^Cre/+^;Rosa26^Ai14/+^* mice. (**I,J**) As in (G,H), but for P21. (**K,L**) As in (G,H), but for P14 *Nlgn2^+/+^;Vip^Cre/+^;Rosa26^Ai14/+^*and *Nlgn2^f/f^*;*Vip^Cre/+^;Rosa26^Ai14/+^*mice. (**M,N**) As in (K,L), but for P21. N number represents # of cells followed by number of animals for cumulative frequency graphs or # of cells for physiology summary data. Each filled (male) or open (female) circle represents one neuron. Bar graphs are mean ± S.E.M. except for sEPSQ:sIPSQ ratio which is median ± interquartile range. *, p<0.05; **, p<0.005. Scale bar is 50 µm.

Based on the high level of Nlgn2 expression in cortical interneurons, we were surprised by the lack of effect on inhibition following Nlgn2 knock-out and sought to determine the efficacy of the interneuron specific Cre lines at deleting Nlgn2. We performed dFISH against *Nlgn2* and *TdTomato* on *Nlgn2^f/f^*;*Sst^Cre/+^;Rosa26^Ai14/+^* and *Nlgn2^+/+^;Sst^Cre/+^;Rosa26^Ai14/+^*controls at P14 and P21 to quantify the levels of *Nlgn2* in Sst interneurons in the presence of *Sst-ires-Cre*. To account for the variability in *Nlgn2* fluorescence between different animals, we normalized the mean fluorescence intensity in Sst cells to the average mean fluorescence intensity of all layer 2/3 cells in the same section. Based on our dFISH comparing *Nlgn2^+/+^* with *Nlgn2^-/-^* animals, 90% of all cells in Nlgn2^-/-^ animals fall below the bottom 10% of cells in the WT animals (Supp Fig 1). Thus, we defined a cell as having *Nlgn2* deleted if the fluorescence intensity was within the bottom 10% of cells in the control. For *Nlgn2^f/f^*;*Sst^Cre/+^;Rosa26^Ai14/+^* at P14 we found only 65.8% of the cells were deleted and at P21 89.7% of the cells were deleted (Fig 1 G-J), indicating a delayed loss of Nlgn2 mRNA. We then performed similar experiments using *Vip-ires-Cre* and *Pv-ires-Cre*. For *Nlgn2^f/f^*;*Vip^Cre/+^;Rosa26^Ai14/^*at P14 we found that 55.0% of the cells were deleted and at P21 77.3% of cells were deleted, again indicating a delayed deletion (Fig 1 K-N).

*Pv-ires-Cre* is known to have delayed expression in cortical interneurons (Taniguchi *et al*., 2011) and our dFISH confirmed this, with no expression of the TdTomato reporter in visual cortex at P14 indicating that Cre was not present (Supp Fig 2 A). Zhang et al., 2016 used *Pv-ires-Cre* to manipulate Nlgn2 in the cerebellum and found a significant decrease in sIPSCs in Purkinje cells but no change in stellate cells despite both cell types expressing Pv-Cre at P21 (Zhang and Südhof, 2016). Our dFISH shows that Pv-Cre efficiently deletes Nlgn2 from Purkinje cells at P14 (Supp Fig 2 B-D). However, when looking at the cerebellar interneurons in the molecular layer, there is clear Cre expression and *Nlgn2* deletion from the presumed basket cells in the inner half of the molecular layer, but no Cre expression in the presumed stellate cells in the outer half of the molecular layer (Supp Fig 2 C). Based on this, we hypothesized that the timing at which Nlgn2 is deleted from a given cell type might be a critical determinant in whether there is an effect on inhibition.

### Early deletion of *Nlgn2* disrupts inhibition onto all three classes of cortical interneurons

In an attempt to achieve earlier deletion of *Nlgn2* in interneurons, we designed a strategy using virally delivery of a Flpo dependent iCre into *Nlgn2 floxed* animals that express Flpo recombinase in different subsets of interneurons. First, we assessed whether this strategy was able to efficiently delete *Nlgn2* by P14. We injected AAV9-FRT-FLEX-iCre-P2A-3xNLS-dTomato into the visual cortex of P0-P2 *Nlgn^f/f^;Rosa26^Ai14/+^;Sst^Flpo/+^*and *Nlgn2^+/+^;Rosa26^Ai14/+^;Sst^Flpo/+^* animals and performed dFISH against *Nlgn2* and *TdTomato* at P14. The Flpo dependent iCre should be reverted to the correct orientation in Flpo+ Sst interneurons, leading to expression of Cre and deletion of *Nlgn2* mRNA (Fig 2 A). We found that 95.4% of Cre+ cells in *Nlgn^f/f^;Rosa26^Ai14/+^;Sst^Flpo/^* ^-^ had deleted Nlgn2 (Fig 2 B,D). Similarly, for *Nlgn^f/f^;Rosa26^Ai14/+^;Vip^Flpo/^* ^-^ we found that 95.0% of cells had deleted *Nlgn2* (Fig 2 C,E). Together, this indicates that viral delivery of iCre achieves a much earlier deletion of Nlgn2 than the interneuron specific Cre lines.

**Figure 2:**
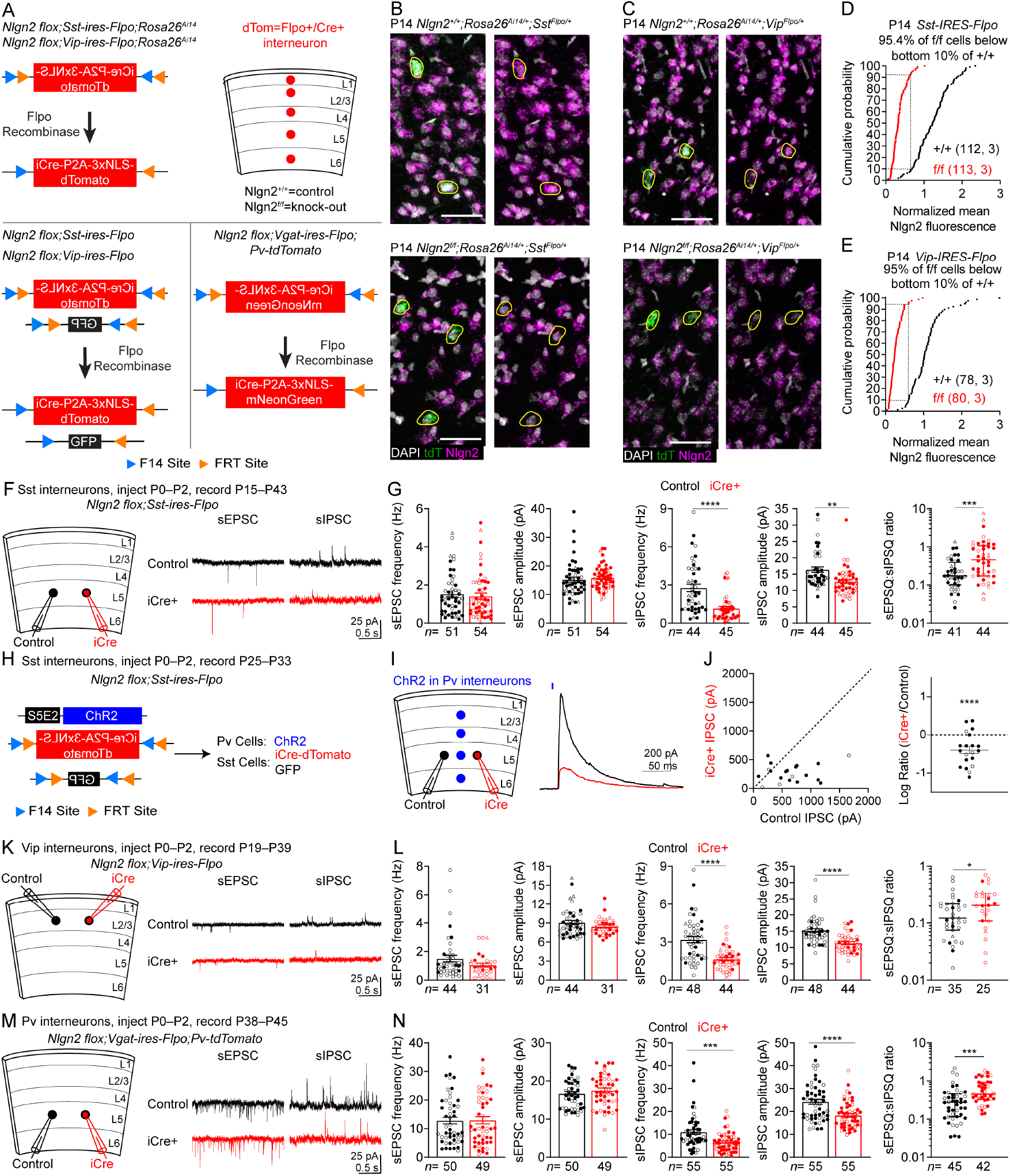
Early deletion of Nlgn2 reduces inhibition onto interneurons. (**A**) Schematic showing viral vectors and experimental strategy. (Top) Genotypes, viral vectors and illustration demonstrating experimental design for dFISH experiments. (Bottom left) Genotypes and viral vectors used for recordings of Sst and Vip positive interneurons. (Bottom right) Genotypes and viral vectors used for recordings of Pv positive interneurons. (**B**) Representative images of DFISH showing *Nlgn2* (magenta), *tdTomato* (green), and DAPI (white) in the visual cortex of P14 *Nlgn2^+/+^;Rosa26^Ai14/+^;Sst^Flpo/+^*and *Nlgn^f/f^;Rosa26^Ai14/+^;Sst^Flpo/+^* mice injected with FFLEX-iCre-P2A-3xNLS-dTomato at P0–P2. Yellow outlines represent *tdTomato* positive cells. (**C**) As in (B), but for *Nlgn2^+/+^;Rosa26^Ai14/+^;Vip^Flpo/+^*and *Nlgn^f/f^;Rosa26^Ai14/+^;Vip^Flpo/+^* mice. (**D**) Cumulative frequencies of tdTomato positive cells as a function of their normalized *Nlgn2* levels. Dashed lines indicate that the Nlgn2 levels of 95.4% of tdTomato positive cells from the P14 *Nlgn2^f/f^*;*Sst^Flpo/+^;Rosa26^Ai14/+^*are within the bottom 10% of tdTomato positive cells from *Nlgn2^+/+^;Sst^Flpo/+^;Rosa26^Ai14/+^*. (**E**) As in (D), but for P14 *Nlgn2^+/+^;Rosa26^Ai14/+^;Vip^Flpo/+^*and *Nlgn^f/f^;Rosa26^Ai14/+^;Vip^Flpo/+^* mice. (**F**) (Left) Schematic illustrating experimental setup for recording of Sst interneurons following early deletion. (Right) Representative traces of sEPSCs (left) and sIPSCs (right) from a control (top) and iCre+ (bottom) Sst cell in layer 5 of visual cortex from *Nlgn2^f/f^;Sst^Flpo/+^* mice injected at P0–P2 with FFLEX-iCre-P2A-dTomato and FFLEX-GFP. (**G**) Summary data of the frequency and amplitude of sEPSCs, sIPSCs, and the sEPSQ:sIPSQ ratio between control and iCre+ cells in *Nlgn2^f/f^;Sst^Flpo/+^* mice following P0–P2 injection. (**H**) Schematic showing viral vectors and genotypes for recording Pv evoked IPSCs onto control and iCre+ Sst interneurons. (**I**) (Left) Schematic illustrating experimental setup for recording Pv evoked IPSCs onto control and iCre+ Sst interneurons. (Right) Representative trace of Pv evoked IPSC onto a control (black) or iCre+ (red) Sst interneuron from layer 5 of visual cortex from *Nlgn2^f/f^;Sst^Flpo/+^* mice injected at P0–P2 with FFLEX-iCre-P2A-3xNLS-dTomato, FFLEX-GFP, and S5E2-ChR2. (**J**) (Left) Summary data of the amplitude of Pv evoked IPSCs onto pairs of control and iCre+ Sst cells from *Nlgn2^f/f^;Sst^Flpo/+^* mice injected at P0–P2. (Right) Summary data of the log ratio of the amplitude of Pv evoked IPSCs onto pairs of control and iCre+ Sst cells *Nlgn2^f/f^;Sst^Flpo/+^* mice injected at P0–P2. (**K,L**) As in (F,G) but for Vip cells in layer 2/3 from *Nlgn2^f/f^;Vip^Flpo/+^* mice. (**M,N**) As in (F,G) but for Pv cells in layer 5 from *Nlgn2^f/f^;Vgat^Flpo/+^;Pv^TdT/+^* mice. N number represents # of cells followed by number of animals for cumulative frequency graphs or # of cells for physiology summary data. Each filled (male) or open (female) circle represents one neuron. Open triangle indicates sex unknown. Bar graphs are mean ± S.E.M. except for sEPSQ:sIPSQ ratio which is median ± interquartile range. *, p<0.05; **, p<0.005; ***, p<0.0005; ****, p<0.0001. Scale bar is 50 µm.

We slightly modified the viral strategy designed above to allow us to identify and record from both *Nlgn2* positive cells and iCre+ *Nlgn2* knock-out cells. To do this, we combined the Flpo dependent iCre virus with a Flpo dependent GFP virus, using a low titer of the Flpo dependent iCre virus and a high titer of the Flpo dependent GFP virus (Fig 2 A). We first determined the titer of AAV9-FRT-FLEX-GFP and AAV9-FRT-FLEX-iCre-P2A-3xNLS-dTomato needed to express GFP in greater than 50% of Sst or Vip neurons and Cre in approximately 10-20% of Sst or Vip interneurons. We estimated the density of Sst, Vip, and Pv cells in visual cortex where we inject the virus by using either *Sst^Flpo/+^;Rosa26^FSF-tdT^* mice for Sst cells (Supp Fig 3 A,D), *Vip^Cre/+^;Rosa26^Ai14/+^* mice for Vip cells (Supp Fig 3 B,D), or *Pv^tdT/+^* mice for Pv cells (Supp Fig 3 C,D). We cut 300 µm live sections from these mice and quantified the density number of tdTomato positive interneurons. We them cut 300 µm live sections from *Nlgn2f/f;SstFlpo/+* (Supp Fig 4 E) or *Nlgn2f/f;VipFlpo/+* (Supp Fig 3 F) animals injected at P0-P2 with AAV9-FRT-FLEX-iCre-P2A-3xNLS-dTomato and AAV9-FRT-FLEX-GFP. We quantified the number of iCre-dTomato and GFP positive cells and estimated the percentage of Sst or Vip cells infected with each virus using the average number of Sst or Vip cells/mm^2^ we calculated (Supp Fig 3 H,I).

To test the effect of early *Nlgn2* deletion in Sst interneurons we cut acute slices from 2-6 week old *Nlgn2^f/f^;Sst^Flpo/+^* animals injected in visual cortex at P0-P2 with AAV9-FRT-FLEX-iCre-P2A-3xNLS-dTomato and AAV9-FRT-FLEX-GFP and recorded sEPSCs and sIPSCs from control and iCre+ Sst neurons in layer 5 (Fig 2 F). We observed no change in sEPSCs but a 60.6% reduction in sIPSC frequency and 22.2% reduction in sIPSC amplitude (Fig 2 G). We verified that decreased inhibition was not due to the presence of virus alone, by performing similar experiments on *Nlgn2^+/+^;Sst^Flpo/+^* animals and found no difference in sEPSCs or sIPSCs (Supp Fig 4 A,B) We next wanted to test whether early *Nlgn2* deletion in Sst interneurons has any effect on evoked inhibition. To do this we used Flpo dependent iCre and Flpo dependent GFP to target the Sst interneurons. We then designed a virus that uses the Pv-specific S5E2 enhancer (Vormstein-Schneider *et al*., 2020) to drive expression of channelrhodopsin (ChR2) specifically in Pv interneurons (Fig 2H). We then simultaneously recorded from a control and *Nlgn2* knock-out Sst interneuron and shined blue light to activate ChR2 and compared the amplitude of the evoked IPSC (Fig 2 I). We found that the amplitude of evoked IPSC in Nlgn2 knock-out Sst interneurons was significantly reduced compared to control Sst interneurons (Fig 2 J)The majority of the Sst cells displayed a small inward photocurrent (mean=70 pA) however this current was much smaller than the evoked IPSC and was present in both control and *Nlgn2* knock-out cells. We hypothesize that this is likely due to a low level of non-specific ChR2 expression, however, this photocurrent was much smaller than the evoked IPSC and was present in both control and *Nlgn2* knock-out cells so it does not affect the interpretation of the data.

To test the effect of early Nlgn2 deletion in Vip neurons, we injected AAV9-FRT-FLEX-iCre-P2A-3xNLS-dTomato and AAV9-FRT-FLEX-GFP into *Nlgn2^f/f^;Vip^Flpo/+^* animals at P0-P2 and recorded sEPSCs and sIPSCs from layer 2/3 Vip neurons (Fig 2 K). We observed no change in sEPSCs but a 47.9% reduction in sIPSC frequency and 25.1% reduction in sIPSC amplitude (Fig 2 L). We verified that decreased inhibition was not due to the presence of virus alone, by performing similar experiments on *Nlgn2^+/+^;Vip^Flpo/+^* animals and found no difference in sEPSCs or sIPSCs (Supp Fig 3 C,D).

We then tested whether using viral delivery of Flpo dependent iCre into *Nlgn2^f/f^;Pv^Flpo/+^* animals had any effect. We injected AAV9-FRT-FLEX-iCre-P2A-3xNLS-dTomato and AAV9-FRT-FLEX-GFP into visual cortex of P0-P2 *Nlgn2^f/f^;Pv^Flpo/+^* animals, and saw no effect on sEPSCS or sIPSCs (Supp Fig 5 A,B). As discussed above (Supp Fig 2) Pv mediated gene expression does not come on until P21, so we hypothesize that the lack of effect on inhibition observed with this experiment is due to the delayed deletion of *Nlgn2*. To test this hypothesis, we modified the experimental strategy to achieve an earlier deletion of Nlgn2 in Pv interneurons. We used *Nlgn2^f/f^;Vgat^Flpo/+^;Pv^TdT/+^*which express Flpo recombinase in all GABAergic interneurons and TdTomato in all Pv interneurons. We then injected a very low titer of AAV9-FRT-FLEX-iCre-P2A-3xNLS-mNeonGreen into these animals to express Cre and delete *Nlgn2* from 10-20% of all interneurons. We estimated the percentage of Pv interneurons expressing iCre by quantifying the number of tdT+ mNeonGreen+ cells and comparing that to the total number of tdT+ cells (Supp Fig 3 G,I). Since Pv interneurons in these mice express TdTomato we were able to identify TdTomato+ control Pv interneurons and TdTomato/mNeonGreen double positive *Nlgn2* knock-out Pv interneurons (Fig 2A). We cut acute slices from 5-7 week old *Nlgn2^f/f^;Vgat^Flpo/+^;Pv-TdT^+/-^*animals injected at P0-P2 with AAV9-FRT-FLEX-iCre-P2A-3xNLS-mNeonGreen and recorded sEPSCs and sIPSCs from Pv neurons in layer 5 (Fig 2 M). We found no change in sEPSCs but a 40.00% reduction in sIPSC frequency and a 24.5% reduction in sIPSC amplitude (Fig 2 N). The difference between interneuron specific Cre line mediated deletion and viral iCre mediated deletion strongly suggests that early deletion of Nlgn2 is required to see effects on inhibition.

### Adult deletion of *Nlgn2* in cortical interneurons has no effect on inhibition

To confirm the timing-dependence of Nlgn2 knock-out in interneurons we modified the above experiment, changing only the age at which the injections were performed. Liang et al. used *Nlgn2 floxed* mice and injected Cre into adult animals. They showed that 2-3 weeks after injection, Nlgn2 protein is gone, indicating that adult deletion is capable of removing protein (Liang *et al*., 2015). We chose to wait six weeks after injection to ensure that all Nlgn2 would be deleted. We cut acute slices from 10 week old *Nlgn2^f/f^;Sst^Flpo/+^* animals that were injected in visual cortex at P30 with AAV9-FRT-FLEX-iCre-P2A-3xNLS-dTomato and FRT-FLEX-GFP and recorded sEPSCs and sIPSCs from control and iCre+ layer 5 Sst neurons, and found no change in either sEPSCs or sIPSCs. (Fig 3 A,B). *For* Vip neurons we performed a similar experiment, recording from layer 2/3 Vip neurons, and again found no change in sEPSCs or sIPSCs (Fig 3 C,D). To confirm that the lack of change in inhibition was due to the change in injection timing and not the older age of the animals, we cut acute slices from 10 week old *Nlgn2^f/f^;Sst^Flpo/+^* animals that were injected at P1 (instead of P30) and found a 60.8% reduction in sIPSC frequency and a trend towards reduced sIPSC amplitude (Supp Fig 8 A,B). Based on the timing-dependence of conditional Nlgn2 deletion phenotypes in interneurons, we next asked if the same thing is true in pyramidal cells.

**Figure 3:**
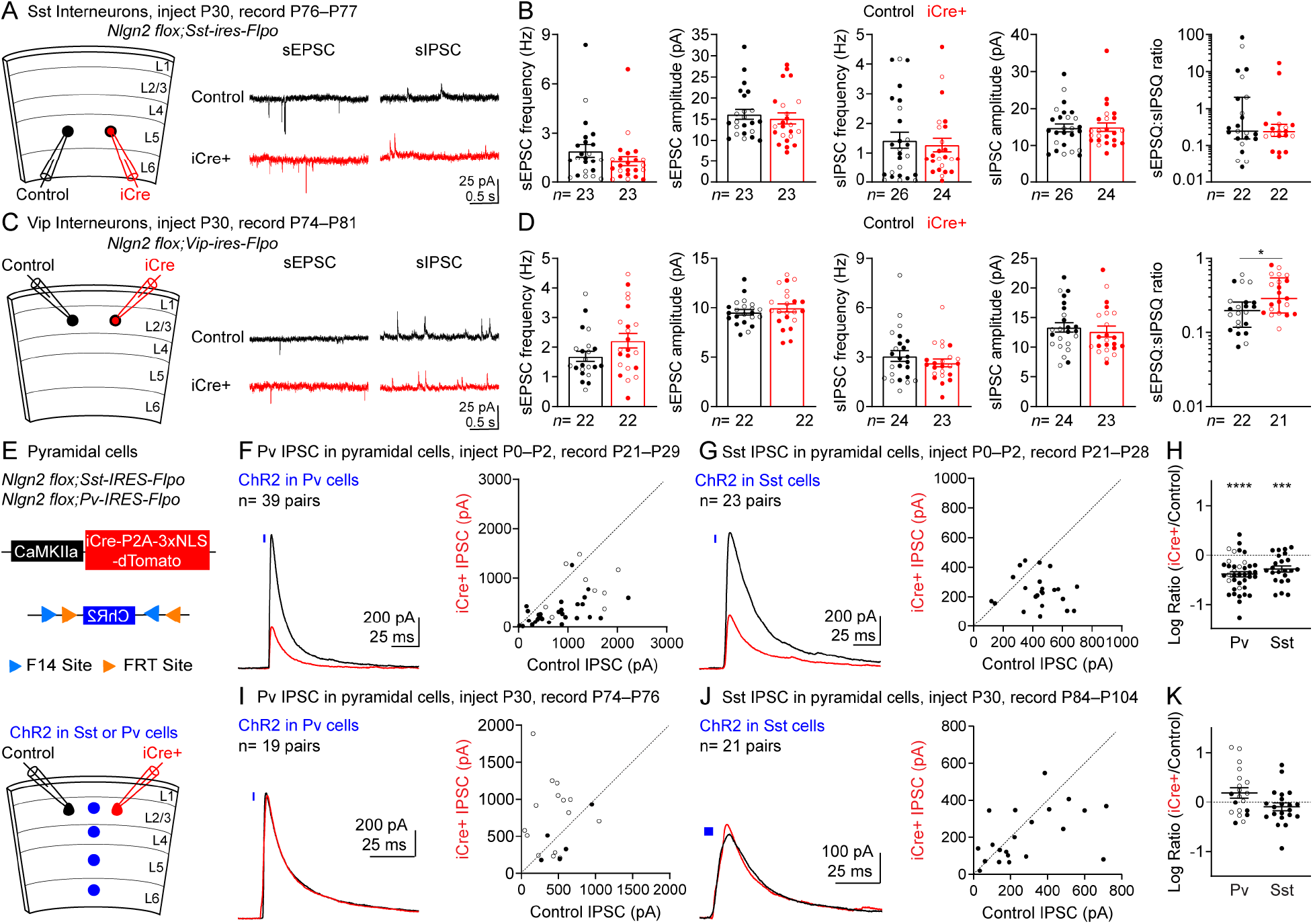
Adult deletion of Nlgn2 has no effect on inhibition onto interneurons or pyramidal cells. (**A**) (Left) Schematic illustrating experimental setup for recording of Sst interneurons following adult deletion. (Right) Representative traces of sEPSCs (left) and sIPSCs (right) from a control (top) and iCre+ (bottom) Sst cell in layer 5 of visual cortex from *Nlgn2^f/f^;Sst^Flpo/+^* mice injected at P30 with FFLEX-iCre-P2A-3xNLS-dTomato and FFLEX-GFP. (**B**) Summary data of the frequency and amplitude of sEPSCs, sIPSCs, and the sEPSQ:sIPSQ ratio between control and iCre+ cells in in *Nlgn2^f/f^;Sst^Flpo/+^* mice following P30 injection. (**C,D**) As in (A,B), but for Vip cells in layer 2/3 from *Nlgn2^f/f^;Vip^Flpo/+^* mice. (**E**) Schematic illustrating genotypes and viral vectors used for recordings of Pv and Sst evoked IPSCs onto pyramidal cells (top) and experimental strategy (bottom). (**F**) (Left) Representative trace of Pv evoked IPSC onto a control (black) or iCre+ (red) layer 2/3 pyramidal cell from *Nlgn2^f/f^;Pv^Flpo/+^* mice following P0–P2 injection of CaMKIIa-iCre-P2A-3xNLS-dTomato and FFLEX-ChR2. (Right) Summary data of the amplitude of Pv evoked IPSCs onto pairs of control and iCre+ layer 2/3 pyramidal cells from *Nlgn2^f/f^;Pv^Flpo/+^*mice following P0–P2 injection. (**G**) As in (F) but for Sst evoked IPSCs from *Nlgn2^f/f^;Sst^Flpo/+^* mice. (**H**) Summary data of the log ratio of the amplitude of Pv and Sst evoked IPSCs onto pairs of control and iCre+ layer 2/3 pyramidal cells. (**I-K**) As in (F-H), but following P30 injection of CaMKIIa-iCre-P2A-3xNLS-dTomato and FFLEX-ChR2. Each filled (male) or open (female) circle represents one neuron. Bar graphs are mean ± S.E.M. except for sEPSQ:sIPSQ ratio which is median ± interquartile range. *, p<0.05; ***, p<0.0005; ****, p<0.0001.

### Early deletion of *Nlgn2* in pyramidal cells reduces inhibition but adult deletion has no effect

We next wanted to test how the effect of sparse deletion of *Nlgn2* in pyramidal cells is affected by the age at which deletion is performed. To sparsely manipulate *Nlgn2* in pyramidal cells we modified the viral delivery of iCre by utilizing a CaMKIIa promoter to drive expression of iCre and dTomato in pyramidal cells. We delivered a low titer of this virus such that only about 5% of all pyramidal cells were labeled. We drove ChR2 expression in either Pv or Sst interneurons to test the effect of *Nlgn2* deletion on evoked IPSCs from both cell types. To do this, we utilized a virus expressing Flpo dependent ChR2 and injected into animals expressing either *Pv-IRES-Flpo* or *Sst-IRES-Flpo*. (Fig 3E). We first wanted to confirm that our viral strategy can efficiently delete *Nlgn2* mRNA at a young age. We performed P0-P2 injection of AAV9-CaMKIIa-iCre-P2A-3xNLS-dTomato into *Nlgn2^f/f^;Rosa26^Ai14/+^*and *Nlgn2^+/+^;Rosa26^Ai14/+^* animals and performed dFISH against *Nlgn2* and *tdTomato* at P18-P21 (Supp Fig 6 A). We found that 95.9% of TdTomato positive cells in *Nlgn2^f/f^;Rosa26^Ai14/+^* animals had Nlgn2 deleted (Supp Fig 6 B,C). We then cut acute slices from 3-4 week old *Nlgn2^f/f^;Sst^Flpo/+^*or *Nlgn2^f/f^;Pv^Flpo/+^* animals injected in visual cortex at P1 with AAV9-CaMKIIa-iCre-P2A-3xNLS-dTomato and AAV9-fDIO-ChR2. We recorded sEPSCs and sIPSCs as well as ChR2 evoked IPSCs from control and iCre+ pyramidal neurons in layer 2/3 (Fig 3 E). There was no change in sEPSCs and a 15.3% reduction in sIPSC frequency and 24.3% reduction in sIPSC amplitude (Supp Fig 6 D,E). Sst and Pv evoked IPSCs were significantly smaller in iCre+ pyramidal cells when compared to nearby WT pyramidal cells (Fig 3 F-H). We confirmed that these effects were due to *Nlgn2* deletion and not just the presence of virus by performing the same experiments on *Nlgn2^+/+^;Sst^Flpo/+^*or *Nlgn2^+/+^;Pv^Flpo/+^* animals. We found no difference in the frequency or amplitude sEPSCs or sIPSCs between control and iCre+ pyramidal cells (Supp Fig 6 F,G). Additionally, we saw no change in the amplitude of Pv (Supp Fig 6 H,J) or Sst (Supp Fig 6 I,J) evoked IPSCs between control and iCre+ cells.

A previous study tested the effect of *Nlgn2* knock-out on Sst and Pv evoked inputs onto pyramidal cells using *Nlgn2^-/-^* animals. They demonstrated that *Nlgn2^-/-^* animals showed a decrease in the amplitude of Pv evoked inputs but no change in the amplitude of Sst evoked inputs comparted to *Nlgn2^+/+^* animals (Gibson, Huber and Südhof, 2009). Interestingly, our sparse deletion of *Nlgn2* showed a decrease in both Sst and Pv evoked synaptic input. Gibson et al. recorded unitary IPSCs by directly stimulating a layer 2/3 Sst interneuron and recording from a nearby pyramidal cell. In our study, ChR2 activation of Sst cells activates a much larger population of cells across all cortical layers. To test whether this might account for the difference between our results we designed a strategy to record unitary IPSCs from Sst interneurons in our sparse deletion model. We generated *Nlgn2^f/f^;Sst^Flpo/+^;Rosa26^FSF-TdTomato/+^* animals which express TdTomato in Sst cells allowing for their identification. We then injected of AAV9-CaMKIIa-iCre-P2A-mNeonGreen to delete *Nlgn2* from about 5% of pyramidal cells while labeling them with mNeonGreen. This allowed for simultaneous recording from an Sst cell, a control pyramidal cell, and an iCre+ pyramidal. We stimulated the Sst cell multiple times to evoke six consecutive action potential while recording from the control and iCre+ pyramidal cell (Supp Fig 6 A). We found no change connection probability between Sst cells and pyramidal cells indicating that *Nlgn2* does not affect the likelihood of synaptic connections between these cell types (Supp Fig 7 B). However, we did find a significant decrease in the amplitude of the first IPSC evoked by the chain of action potentials in iCre+ cells compared to control cells (Supp Fig 7 C). We then normalized the amplitude of each IPSC in the spike train for a given cell to the first IPSC for that cell to compare spike train potentiation. We found no difference between control and iCre+ cells (Supp Fig 7 C). Our finding that uIPSCs between Sst cells and *Nlgn2* knock-out pyramidal cells are altered in our sparse deletion model suggests that the difference between our findings and those of Gibson et al. are likely due to the method of deleting *Nlgn2* (sparse cell-type specific vs global knock-out) or the cortical region tested. More follow-up studies need to be performed to address these possibilities more thoroughly.

After confirming that early sparse viral deletion of Nlgn2 in pyramidal cells decreases inhibition, like our observations in interneurons, we next asked whether these effects are also dependent on the timing of *Nlgn2* deletion. We performed the same experiment described above (Fig 3 E) but modified it by changing the timing of the viral injection from P0-P2 to P30 and then waiting 6 weeks following viral injection. In this case, we observed no change in sEPSCs or sIPSC frequency and a 14.3% decrease in sIPSC amplitude (Supp Fig 6 K,L) Despite the small effect on sIPSC amplitude, we observed no change in Pv or Sst evoked IPSCs between control and iCre+ cells (Fig 3 I-K). We then tested whether the small effect on sIPSC amplitude was due to viral expression by performing the same experiment in *Nlgn2^+/+^* animals. In this case we found no difference in sEPSCs or sIPSCs suggesting viral expression alone does not lead to changes in inhibition (Supp Fig 6 M,N). In this experiment, we performed our recording on animals that were ∼10 weeks old versus the 3–4-week-old animals we tested following P0-P2 injection. To test the older age of animals did not contribute to the smaller effect on inhibition observed, we performed viral injection at P0-P2 and waited until the animals were 10 weeks of age to record sPSCs and Sst evoked IPSCs. We found a 32.5% reduction in sIPSC frequency and a 27.3% reduction in sIPSC amplitude (Supp Fig 8 C,D), and a significant decrease in the amplitude of Sst evoked IPSCs between WT and iCre+ cells (Supp Fig 8 E,F).

Another study conditionally deleted *Nlgn2* in the prefrontal cortex in adult animals and showed that after 2-3 weeks there were behavioral changes consistent with *Nlgn2* deletion and after 6-7 weeks there was a significant decrease in the frequency and amplitude of sIPSCs (Liang *et al*., 2015). While Liang et al. deleted *Nlgn2* from all neurons in the prefrontal cortex using a high titer of a virus driving Cre expression with the human synapsin promoter, our study performed a sparse deletion, specifically studying the cell-autonomous effects of deletion. We hypothesized that this might explain the lack of effect we observed in our study. To test this, we injected a high titer of AAV9-hSyn-eGFP-Cre virus into *Nlgn2^f/f^* animals to delete *Nlgn2* from all neurons in the local area of our injection in visual cortex. As a control, we performed the same viral injection into *Nlgn2^+/+^*animals (Supp Fig 8 G). We found no difference in sEPSCs or sIPSC amplitude but we observed a significant reduction in sIPSC frequency (Supp Fig 8 H,I). This suggests that cell-autonomous deletion of *Nlgn2* from pyramidal cells has no effect on sIPSC frequency while large scale deletion of *Nlgn2* disrupts the cortical network such that sIPSC frequency is altered. Additional follow-up studies are needed to more specifically investigate the differences between cell-autonomous *Nlgn2* deletion and more global *Nlgn2* deletion in pyramidal cells.

Based on our data showing that early Nlgn2 deletion in both excitatory and inhibitory cells disrupt inhibition, we next asked how specifically disrupting inhibition onto excitatory cells or onto inhibitory cells might alter E-I balance and cortical function.

### Conditional *Nlgn2* deletion in excitatory neurons is lethal but deletion in both excitatory and inhibitory neurons simultaneously partially restores viability

To understand how inhibitory synapses onto excitatory neurons and inhibitory synapses onto inhibitory neurons contribute to E-I balance we conditionally deleted *Nlgn2* in excitatory and inhibitory neurons. First we attempted to breed *Nlgn2^f/f^;Vglut2^Cre/+^*(Vglut2-cKO) animals to study how disrupting inhibitory synapses onto excitatory neurons while preserving inhibitory synapses onto interneurons impacts E-I balance. To our surprise *the* behavioral impacts of conditional Nlgn2 deletion in excitatory neurons we bred Nlgn2 floxed mice with Vglut2-Cre mice to remove Nlgn2 from all glutamatergic neurons. Much to our surprise, we found that Vglut2-cKO leads to complete lethality. Based on our breeding scheme we would have expected to observe 22 Vglut2-cKO animals however we did not observe a single one (Fig 4 A). We next attempted to breed *Nlgn2^f/f^;Vgat^Cre/+^*(Vgat-cKO) animals to study how disrupting inhibitory synapses onto interneurons while preserving those onto excitatory neurons impacts E-I balance. Again, we found that Vgat-cKO displayed significant lethality. Out of 22 expected Vgat-cKO animals, we observed 3, for a 12.6% survival (Fig 4 B).

**Figure 4:**
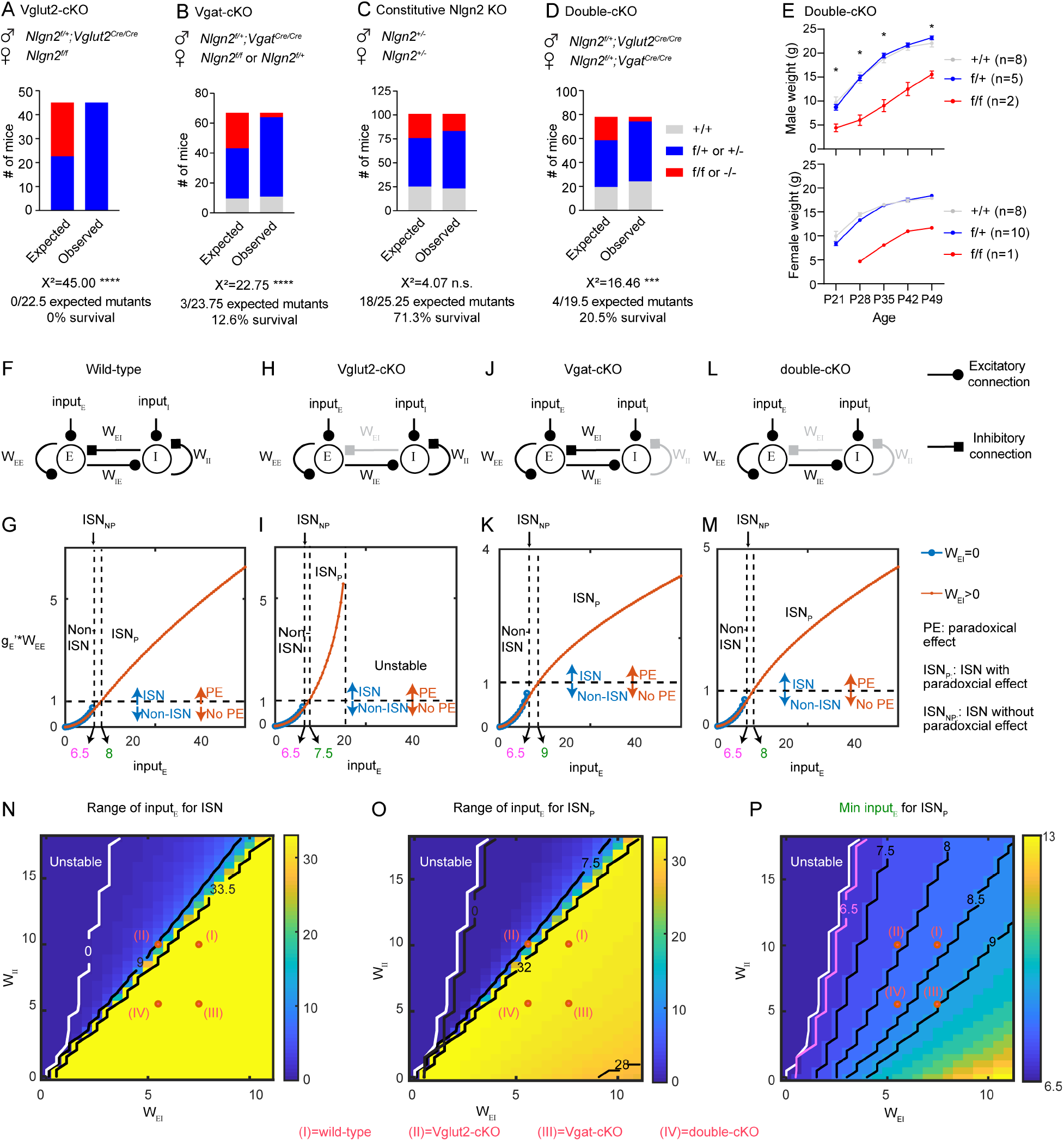
Conditional deletion of Nlgn2 in excitatory or inhibitory neurons is lethal, potentially due to abnormal network dynamics. (**A**) Survival data for *Nlgn2 floxed;Vglut2-Cre* conditional knock-out of Nlgn2 in excitatory neurons. Χ^2^ value compares observed offspring to expected Mendellian ratio for the depicted breeding scheme. (**B**) As in (A), but for *Nlgn2 floxed; Vgat-Cre* conditional knock-out of Nlgn2 in inhibitory neurons. (**C**) As in (A), but for constitutive *Nlgn2* knock-out in all cells. (**D**) As in (A), but for *Nlgn2 floxed; Vglut2-Cre; Vgat-Cre* double conditional knock-out of Nlgn2 in excitatory and inhibitory neurons. (**E**) Summary data of the body weight for male and female double-cKO mice and controls from 3–7 weeks of age. Stars indicate results of two-way ANOVA with Tukey’s multiple comparison test between *Nlgn2^+/+^* and *Nlgn2^f/f^*. N number represents # of mice. n.s., p>0.05; **, p<0.005; ****, p<0.0001. (**F**)-(**J**) A computational model of an E-I network, which operates in different dynamical regimes depending on the external input to E-population (*input_E_*). (**F**) Schematic of the computational model corresponding to four experimental scenarios explored in this paper (see **Methods** for additional details). The effect of conditional deletion of Nlgn2 is mimicked by weakening the strength of the inhibitory connections (grey) onto specific types of cells. (**F1**) Wildtype. (**F2**) Vglut2-cKO, reduced *W_EI_*. (**F3**) Vgat-cKO, reduced *W_II_*. (**F4**) Double-cKO, reduced *W_EI_* and *W_II_*. (**G**) The operating regime of the network changes with the external input *input_E_* in a connectivity-dependent manner. Blue curve: value of 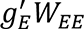 at the stationary state for the network without feedback inhibition (*W_IE_* = 0) as a function of the external input *input_E_*. The network without feedback inhibition is stable if and only if this term is smaller than 1. Red curve: same as the blue curve but for the network with feedback inhibition (*W_IE_* > 0). The network exhibits a paradoxical effect if and only if this term is larger than 1. With increasing *input_E_*, the blue curve vanishes, which implies a transition from non-ISN regime to ISN regime. With sufficiently large *input_E_*, the paradoxical effect emerges (from ISN_NP_ to ISN_P_) as the red curve crosses 1. In (**G2**), with sufficiently large *input_E_*, the red curve vanished, indicating the instability of the network with feedback inhibition. The input thresholds for the transition from non-ISN to ISN (magenta) and the emergence of paradoxical effect (green) are labeled. Note that the input threshold for an ISN regime is independent of *W_EI_* and *W_II_*. Amplitude of reduction in *W_EI_* and *W_II_* compared to the scenario of wild type: (I) 0%, 0%. (II) 26.7%, 0%. (III) 0%, 45%. (IV) 26.7%, 45%. (**H**) The range of *input_E_* allowing an ISN operating regime as a function of the strength of the inhibitory connections onto excitatory cells (*W_EI_*) and inhibitory cells (*W_II_*). Under the parameter regime on the left of the contour line at 0 (white), the network can never operate in an ISN regime since it loses stability from non-ISN regime with increasing *input_E_*. Same area is marked by white in (**I**) and (**J**). On the right of the contour line at 33.5, the network operates in an ISN regime under the maximal value of *input_E_* (*max*(*input_E_*) = 40) we explored. (**I**) The range of *input_E_* allowing a paradoxical effect as a function of *W_EI_* and *W_II_*. (**J**) The threshold of *input_E_* to obtain a paradoxical effect as a function of *W_EI_* and *W_II_*. In (**H**)-(**J**), the parameters for four examples shown in (**G**) are marked.

The lethality we observed in Vglut2-cKO and Vgat-cKO animals is very surprising because constitutive Nlgn2 deletion is known to be viable (Blundell *et al*., 2009; Gibson, Huber and Südhof, 2009; Poulopoulos *et al*., 2009). Based on our findings, we sought to determine whether there might be subtle survival defects in *Nlgn2^-/-^*animals. We analyzed our breeding data and found no significant change in survival for constitutive Nlgn2 knock-out animals. Out of 25 expected mutant animals we observed 18, for a 71.3% survival. This slight decrease in survival was not significant (Fig 4 C).

To further understand why constitutive Nlgn2 knock-out is viable and Vglut2-cKO and Vgat-cKO are lethal we attempted to generate *Nlgn2^f/f^;Vglut2^Cre/+^;Vgat^Cre/+^* (double-cKO) animals to simultaneous disrupt inhibitory synapses onto excitatory and inhibitory neurons. Out of 19 expected double-cKO animals we observed 4, for 20.5% survival (Fig 4 D). The four surviving animals that we observed had significantly decreased body weight at all ages compared to their control littermates (Fig 4 E), suggesting that they might display developmental delay similar to what has been shown for *Nlgn2^-/-^* animals (Wöhr *et al*., 2013). While the double-cKO animals still displayed significant lethality, the survival was significantly better than what was observed for Vglut2-cKO animals, suggesting that removing Nlgn2 in inhibitory cells as well as excitatory cells can somewhat restore survival. We observed a small increase in survival in double-cKO animals compared to Vgat-cKO animals suggesting that removing Nlgn2 in excitatory cells might improve survival compared to removal in inhibitory cells alone, however more animals need to be studied to confirm this finding. Interestingly double-cKO animals display much worse survival compared to constitutive knock-out animals suggesting that other cell types might be contributing. This will be interesting to address in future studies.

To our knowledge, there is no other gene that has been shown to have such a pronounced lethality phenotype following cell-type specific deletion and no lethality at all following complete loss of the gene. We hypothesize that this might be due to the manipulation of inhibitory synapses onto one type of cells without affecting the others. This would suggest that inhibitory synapses onto both excitatory and inhibitory cells are critical for E-I balance and cortical function. Unfortunately, due to the lethality of Vglut2-cKO and Vgat-cKO animals, it is not possible to investigate E-I balance in these animals experimentally. To address this question, we turned to computational modeling of a simple cortical network.

### A computational model demonstrates the impact of the deletion of *Nlgn2* on network dynamics

To gain insights into the impact of the deletion of Nlgn2 and consequential altered E-I balance at the network level, we developed a computational model using a canonical E-I network. This model builds on a previous inhibition-stabilized network (ISN) model with a supralinear response function. The model involves a single population of inhibitory cells and a population of excitatory cells (Fig 4F). The state of the network is characterized by the population firing rates of E cells and I cells. The nonlinearity of the transfer function enables the network to transition between two different operating dynamical regimes: a non-ISN regime for weak external input (*input_E_*) and an ISN regime for strong feedforward drive (*input_E_*), in which the paradoxical effect arises. Such an input-dependent behavior empowers the model to explain various cortical computations and properties, including multi-input integration and contextual modulation. In our model, the effect of conditional deletion of Nlgn2 was mimicked by weakening the strength of the inhibitory connections (Fig 4F) onto specific types of cells. In this part, we focused on the impact of the deletion of Nlgn2 on the network dynamics, paradoxical effect, and their dependence on the feedforward input. We approached that by numerically investigating the operating dynamical regime of the network with varying levels of external input to the excitatory population for four scenarios shown in Fig 4 F.

The operating dynamical regime and the emergence of paradoxical effects under varying levels of external input were assessed by spectral analysis on the network connectivity at the stationary state (see Methods). Note that this analysis could be done numerically only when the corresponding network is stable. For the wild-type scenario (Fig G1), with increasing external input, the network transitioned from a non-ISN regime to an ISN-regime. Because of the supra-linearity of the response function, the threshold of input for the emergence of paradoxical effect (green) was higher than that for the transition of the operating dynamical regime (magenta). Note that the latter was not influenced by the inhibitory connection strength (see Methods) and therefore was uniform for four explored scenarios. For scenarios of Vgat-cKO and double-cKO, networks displayed qualitatively similar input-dependent behavior (Fig 4 G3, 4). The most prominent effect of the deletion of Nlgn2 to the network dynamics was found in the scenario of Vglut2-cKO (Fig 4 G2): although the network displayed a similar input-dependent behavior as the wild-type network in a lower range of input, it lost stability (vanished red curve in Fig 4 G2) after it entered ISN regime and displayed paradoxical effect with increasing input, suggesting that the network might not be functioning properly. Such instability was caused by the attenuation in the strength of the inhibitory connections onto excitatory cells (*W_EI_*) and the consequential failure for the inhibitory feedback loop within the network to balance the excitation. This instability also leads to a shorter range of *input_E_* allowing ISN regime (Fig 4H). Surprisingly, this instability and the induced shrinking of input range allowing ISN regime could be rescued by strengthening the inhibitory feedback loop via additionally attenuating the strength of the disinhibitory connection onto inhibitory cells (*W_II_*), i.e., double-cKO (Fig 4 G4, H).

The paradoxical effect, in which increasing direct excitatory input to inhibitory cells leads to a paradoxical decrease in their activity, arises when the network dynamics is dominated by the recurrent, within-network inhibition rather than feedforward drive. It has been observed in many brain regions and proposed to explain multiple cortical properties such as surround suppression. Motivated by that, we addressed the question of how the deletion of Nlgn2 impacted the emergence of paradoxical effects. To do that, across a range of values for the strengths of the inhibitory connection onto excitatory (*W_EI_*) and inhibitory cells (*W_II_*), we determined the range and the threshold of external input (*input_E_*) giving rise to paradoxical effect. As shown in Fig 4 I, due to the instability caused by the attenuated inhibitory feedback loop, networks with a weaker (*W_EI_*) or a stronger *W_II_* showed a significantly shorter range with paradoxical effect. The threshold of external input for paradoxical effect increased with *W_EI_* and decreases with *W_II_* (Fig 4 J). Therefore when the external input was not strong enough, Vgat-cKO can lead to the loss of paradoxical effect. Again, both the shorter range of input allowing paradoxical effect induced by Vglut2-cKO and the higher threshold of input for paradoxical effect induced by Vgat-cKO could be rescued by decreasing *W_EI_*, *W_II_* together, i.e., double-cKO. As marked in Fig 4I, J, the networks investigated in Fig 4F, G for scenarios of wild-type and double-cKO show similar performance.

Overall, our cortical network model suggests that decreasing one subtype of inhibition without proportionally decreasing the other leads to irregularities in performance of the cortical network including increased instability and altering the threshold of external input at which the network is able to function as an ISNP. However, both irregularities can be rescued by double-cKO. While it is difficult to recapitulate lethality in a model of a cortical network, the results reflect similar trends as our observations of lethality in conditional Nlgn2 knock-out animals. Future studies are needed to understand the implications of the model predictions and their relationship to the experimental data.

## Discussion

E-I balance is one of the foundational concepts underlying cortical function. Whether the brain is performing simple computations like processing sensory stimuli, or the more complex computations required for expressing emotion, there is a proportionality between the amount of excitatory and inhibitory activity. The criticality of E-I balance is further evidenced by its implication in neuropsychiatric diseases, however definitive causal evidence linking E-I balance and ASD is scarce. In this study, we manipulated E-I balance throughout development by selectively deleting *Nlgn2* in either excitatory neurons or inhibitory neurons.

To utilize conditional *Nlgn2* knock-out as a tool to study inhibitory synapses onto interneurons we first had to confirm that *Nlgn2* regulates inhibition in interneurons. To our surprise, using *Sst-ires-Cre*, *Vip-ires-Cre*, or *Pv-ires-Cre* to selectively delete *Nlgn2* in each of the three interneuron subtypes had no effect on their excitation or inhibition. This led us to investigate the efficacy of these Cre lines in deleting *Nlgn2*. We demonstrated that *Sst-ires-Cre* and *Vip-ires-Cre* do not adequately delete *Nlgn2* by P14 however they achieve a much more robust deletion by P21. Meanwhile *Pv-ires-Cre* is not even expressed in cortical interneurons at P14. We then developed a Flpo dependent viral method which achieves earlier cell-type specific of *Nlgn2* in interneurons. Using this method, we showed that sparse deletion of *Nlgn2* in all three subtypes of cortical interneurons reduces their inhibition without disrupting their excitation.

Based on our finding that interneuron specific Cre lines are delayed in their deletion of *Nlgn2* and have no effect on inhibition, we investigated the age dependence of *Nlgn2* deletion. We found that adolescent deletion of *Nlgn2* after the formation of synapses is complete does not disrupt their inhibitory synaptic input. This is true for both interneurons as well as pyramidal cells. This suggests that *Nlgn2* may be playing an important role in the development of inhibitory synapses while it is dispensable for their maintenance. This will be interesting to investigate in future studies by looking at the number of inhibitory synapses onto *Nlgn2* knock-out interneurons as well as the molecular components of those synapses.

Given that *Nlgn2* regulates inhibitory synapses onto both excitatory neurons and inhibitory neurons, it is a viable tool to study the contribution of the two different synaptic subtypes to E-I balance. We showed that selectively reducing inhibitory synaptic input onto excitatory or inhibitory neurons results in significant lethality. This lethality can be partially rescued by simultaneously removing *Nlgn2* from both excitatory and inhibitory neurons and constitutive *Nlgn2* knock-out mice display normal survival, although these mice do have developmental delay and anxiety (Blundell *et al*., 2009). This suggests that manipulating inhibitory synapses onto either inhibitory or excitatory neurons individually causes a very severe disruption in E-I balance that results in lethality while disrupting both types simultaneously causes a more mild disruption of E-I balance, allowing the mice to survive. While Vgat-cKO and Vglut2-cKO mice display lethality, it is possible that disrupting *Nlgn2* in excitatory or inhibitory neurons of the forebrain might result in viable animals where we can more directly study the contribution of these two synaptic subtypes to E-I balance.

While we were unable to experimentally test the E-I balance in the Vgat and Vglut2-cKO mouse models due to their lethality, we utilized computational modeling to examine the effects of selectively disrupting the two sub-types of inhibition. Our model suggests that decreasing inhibition onto excitatory neurons while maintaining inhibition onto inhibitory neurons leads to significant instability of the cortical network at higher levels of sensory input. We hypothesize that this is due to high levels of dis-inhibition of pyramidal cells in addition to their decreased activity leading to unstable levels of excitatory activity. Selectively decreasing inhibition onto inhibitory neurons while maintaining inhibition onto excitatory neurons led to more subtle changes in the range of inputs at which the network functions as an ISN_P_. FThis suggest that there is an abnormality in the range of activity where network dynamics are dominated by recurrent within-network inhibition, and may cause abnormalities in functions like surround suppression. Finally, decreasing inhibition onto both excitatory and inhibitory neurons results in a network that performs similarly to the wild-type network. Our double-cKO mouse model does have significant lethality defects and constitutive knock out mice display behavioral and developmental phenotypes suggesting that their cortical network is still abnormal, however it is clearly milder than the Vgat-cKO or Vglut2-cKO. A more complex model that better represents the different types of neurons and connections in the cortex might better capture these more subtle changes.

## Methods

### Mice

*Nlgn2* flox (JAX #025544), *Sox2-Cre* (JAX #008454), *Sst-ires-Cre* (Jax #013044), *Vip-ires-Cre* (JAX #010908), *Pv-ires-Cre* (JAX #017320), *Sst-ires-Flpo* (JAX #028579), *Vip-ires-Flpo* (JAX #028578), *Pv-2A-Flpo* (Jax #022730), *Viaat-ires-Cre* (JAX #028862), *Vglut2-ires-Cre* (JAX #016963), *Pv-tdTomato* (JAX #027395), *Rosa26-CAG-LSL-tdTomato* (Ai14 line, JAX #007914), and *Rosa26-CAG-FSF-LSL-tdTomato* (Ai65 line, JAX #021875) mice were obtained from the Jackson Laboratory. *Nlgn2* KO mice were generated by crossing *Nlgn2* flox mice with *Sox2-Cre* mice to delete the floxed exons 4–6. *Rosa26-CAG-FSF-tdTomato* mice were generated by crossing *Rosa26-CAG-FSF-LSL-tdTomato* mice with *Sox2-Cre* mice to remove the LSL cassette. *Viaat-ires2-Flpo* (JAX #031331) mice were obtained from Dr. Hongkui Zeng at the Allen Institute for Brain Science. All mice were maintained on the C57BL/6J background except *Sst-ires-Cre, Vip-ires-Cre, Sst-ires-Flpo*, and *Vip-ires-Flpo* mice that were on a C57BL/6;129S4 mixed background. *Nlgn2* KO mice were genotyped by PCR using a common forward primer 5’-CAGGAGCAGGAAGGAGACTTGTG-3’, a reverse primer located downstream of the deleted region 5’-CCTCCACGAACTTGAGACCCT-3’ and a reverse primer located within the deleted region, 5’-CTCCTCATCACTGGCCAGCAT-3’. *Rosa26-CAG-FSF-tdTomato* mice were genotyped by PCR using primer sets 5’-AAGGGAGCTGCAGTGGAGTA-3’ and 5’-CCGAAAATCTGTGGGAAGTC-3’ for WT allele and 5’-GGCATTAAAGCAGCGTATCC-3’ and 5’-CTGTTCCTGTACGGCATGG-3’ for the knockin allele. Both male and female mice were used for all experiments. Mice were housed in an Association for Assessment and Accreditation of Laboratory Animal Care International-certified animal facility on a 14-hour/10-hour light/dark cycle. All procedures to maintain and use mice were approved by the Institutional Animal Care and Use Committee at Baylor College of Medicine (protocol AN-6544).

### DNA constructs

Plasmids pAAV-EF1α-DIO-hChR2(H134R)-mCherry (Addgene #20297), pAAV-CaMKIIα-eArchT3.0-P2A-EGFP (Addgene #51110), pAAV-EF1α-DIO-oChIEF(E163A/T199C)-P2A-dTomato (Addgene #51094), pCAG-EGFP (Addgene #11150), pAAV-EF1α-fDIO-hChR2(H134R)-EYFP (Addgene #55639), and pAAV-S5E2-dTomato-P2A-nls-dTomato (Addgene #135630) were obtained from Addgene, pCAGGS-iCre-Puro from Dr. A. Francis Stewart at Technische Universität Dresden, pAAV-CAG-mNeonGreen (Addgene #99134) from Dr. Viviana Gradinaru at California Institute of Technology, and pCAG-tdTomato from Dr. Anirvan Ghosh at University of California, San Diego. Plasmids pAAV-EF1α-FRT-FLEX-mNaChBac-T2A-tdTomato (Addgene #60658) and pCAG-hChR2(H134R)-EYFP (Addgene #114367) were described previously {Xue:2014dl, Messier:2018fo}. All other plasmids were generated and deposited at Addgene as below. pAAV-EF1α-DIO-tdTomato (Addgene #190767) were generated by replacing the hChR2(H134R)-mCherry in pAAV-EF1α-DIO-hChR2(H134R)-mCherry with the tdTomato from pCAG-tdTomato. pAAV-CaMKIIα-iCre-P2A-3xNLS-dTomato (Addgene #190768) was generated by replacing the eArchT3.0-P2A-EGFP of pAAV-CaMKIIα-eArchT3.0-P2A-EGFP with the iCre-P2A-3xNLS-dTomato, which was generated by PCR amplification from pCAGGS-iCre-Puro and pAAV-EF1α-DIO-oChIEF(E163A/T199C)-P2A-dTomato. pAAV-CaMKIIα-iCre-P2A-mNeonGreen (Addgene #190769) was generated by replacing the 3xNLS-dTomato in pAAV-CaMKIIα-iCre-P2A-3xNLS-dTomato with the mNeonGreen from pAAV-CAG-mNeonGreen. pAAV-EF1α-FRT-FLEX-EGFP (Addgene #190770) was generated by replacing the mNaChBac-T2A-tdTomato of pAAV-EF1α-FRT-FLEX-mNaChBac-T2A-tdTomato with the EGFP from pCAG-EGFP. pAAV-EF1α-FRT-FLEX-MCS (Addgene #190771) was generated by replacing the DIO-oChIEF(E163A/T199C)-P2A-dTomato of pAAV-EF1α-DIO-oChIEF(E163A/T199C)-P2A-dTomato with the FRT-FLEX-MCS of pJ244-FRT-FLEX-MCS {Xue:2014dl}. The iCre-P2A-3xNLS-dTomato from pAAV-CaMKIIα-iCre-P2A-3xNLS-dTomato was then inserted into pAAV-EF1α-FRT-FLEX-MCS to generate pAAV-EF1α-FRT-FLEX-iCre-P2A-3xNLS-dTomato (Addgene #190772). pAAV-EF1α-FRT-FLEX-iCre-P2A-3xNLS-mNeonGreen (Addgene #190773) was generated by replacing the dTomato of pAAV-EF1α-FRT-FLEX-iCre-P2A-3xNLS-dTomato with the mNeonGreen from pAAV-CAG-mNeonGreen. pAAV-S5E2-hChR2(H134R)-EYFP (Addgene #190774) was generated by replacing the dTomato-P2A-nls-dTomato of pAAV-S5E2-dTomato-P2A-nls-dTomato with the hChR2(H134R)-EYFP from pCAG-hChR2(H134R)-EYFP.

### AAV production and injection

All recombinant AAV serotype 9 vectors were produced by the Gene Vector Core at Baylor College of Medicine, except for AAV9-hSyn-HI-EGFP-Cre, which was purchased from Penn Vector Core (PL-C-PV1848). AAV vectors were injected into the primary visual cortices of neonatal mice at postnatal day 0–2 (P0–2) with 100–200 nl of virus or young adult mice at postnatal day 30 (P30) with 200 nl of virus (see below). Injection was performed with an UltraMicroPump III and a Micro4 controller (World Precision Instruments) as previously described {Xue:2014dl, Messier:2018fo}.

To express tdTomato in Sst cells of *Nlgn2^f/f^;Sst^Cre/+^*and *Nlgn2^+/+^;Sst^Cre/+^* mice at P0–2, 200 nl of AAV9-EF1α-DIO-tdTomato (6.1×10^11^ GC/ml) was used. To express iCre-P2A-3xNLS-dTomato and EGFP in Sst cells of *Nlgn2^f/f^;Sst^Flpo/+^*and *Nlgn2^+/+^;Sst^Flpo/+^* mice at P0–2 or P30, a 200 nl mixture of AAV9-EF1α-FRT-FLEX-iCre-P2A-3xNLS-dTomato (4.3×10^10^ GC/ml) and AAV9-EF1α-FRT-FLEX-EGFP (5.5×10^12^ GC/ml) was used. To express iCre-P2A-3xNLS-dTomato and EGFP in Sst cells and ChR2 in Pv cells of *Nlgn2^f/f^;Sst^Flpo/+^*mice at P0–2, a 200 nl mixture of AAV9-EF1α-FRT-FLEX-iCre-P2A-3xNLS-dTomato (4.3×10^10^ GC/ml), AAV9-EF1α-FRT-FLEX-EGFP (5.63×10^11^ GC/ml), and AAV9-S5E2-hChR2(H134R)-EYFP (2.46×10^13^ GC/ml) was used. To express iCre-P2A-3xNLS-dTomato in Sst cells of *Nlgn2^f/f^;Sst^Flpo/+^*;*Rosa26^Ai14/+^* and *Nlgn2^+/+^;Sst^Flpo/+^;Rosa26^Ai14/+^* mice at P1, 200 nl of AAV9-EF1α-FRT-FLEX-iCre-P2A-3xNLS-dTomato (4.3×10^10^ GC/ml) was used. To express iCre-P2A-3xNLS-dTomato and EGFP in Vip cells of *Nlgn2^f/f^;Vip^Flpo/+^* and *Nlgn2^+/+^;Vip^Flpo/+^*mice at P0–2 or P30, a 200 nl mixture of AAV9-EF1α-FRT-FLEX-iCre-P2A-3xNLS-dTomato (8.67×10^10^ GC/ml) and AAV9-EF1α-FRT-FLEX-EGFP (5.5×10^12^ GC/ml) was used. To express iCre-P2A-3xNLS-mNeonGreen in Vip cells of *Nlgn2^f/f^;Vip^Flpo/+^*;*Rosa26^Ai14/+^*and *Nlgn2^+/+^;Vip^Flpo/+^;Rosa26^Ai14/+^* mice at P1, 200 nl of AAV9-EF1α-FRT-FLEX-iCre-P2A-3xNLS-mNeonGreen (2.15×10^11^ GC/ml) was used. To express iCre-P2A-3xNLS-dTomato and EGFP in Pv cells of *Nlgn2^f/f^;Pv^Flpo/+^* mice at P0–2, a 200 nl mixture of AAV9-EF1α-FRT-FLEX-iCre-P2A-3xNLS-dTomato (1.3×10^11^ GC/ml) and AAV9-EF1α-FRT-FLEX-EGFP (5.5×10^12^ GC/ml) was used. To express iCre-P2A-3xNLS-mNeonGreen in Pv cells of *Nlgn2^f/f^;Viaat^Flpo/+^;Pv-tdTomato^Tg/+^*mice at P0–2, 200 nl of AAV9-EF1α-FRT-FLEX-iCre-P2A-3xNLS-mNeonGreen (8.6×10^10^ GC/ml) was used. To express iCre-P2A-3xNLS-dTomato in pyramidal cells and ChR2 in Sst cells of *Nlgn2^f/f^;Sst^Flpo/+^* and *Nlgn2^+/+^;Sst^Flpo/+^*mice at P0–2, a 100 nl mixture of AAV9-CaMKIIα-iCre-P2A-3xNLS-dTomato (1.4×10^10^ GC/ml or 1.05×10^11^ GC/ml) and AAV9-EF1α-fDIO-hChR2(H134R)-EYFP (1.89×10^14^ GC/ml) was used. To express iCre-P2A-3xNLS-dTomato in pyramidal cells and ChR2 in Pv cells of *Nlgn2^f/f^;Pv^Flpo/+^*and *Nlgn2^+/+^;Pv^Flpo/+^* mice at P0–2, a 100 nl mixture of AAV9-CaMKIIα-iCre-P2A-3xNLS-dTomato (2.1×10^10^ GC/ml) and AAV9-EF1α-fDIO-hChR2(H134R)-EYFP (1.89×10^14^ GC/ml) was used. To express iCre-P2A-3xNLS-dTomato in pyramidal cells and ChR2 in Sst cells of *Nlgn2^f/f^;Sst^Flpo/+^*mice at P30, a 200 nl mixture of AAV9-CaMKIIα-iCre-P2A-3xNLS-dTomato (1.9×10^10^ GC/ml) and AAV9-EF1α-fDIO-hChR2(H134R)-EYFP (1.3×10^12^ GC/ml) was used. To express iCre-P2A-3xNLS-dTomato in pyramidal cells and ChR2 in Pv cells of *Nlgn2^f/f^;Pv^Flpo/+^* mice at P30, a 200 nl mixture of AAV9-CaMKIIα-iCre-P2A-3xNLS-dTomato (1.9×10^10^ GC/ml) and AAV9-EF1α-fDIO-hChR2(H134R)-EYFP (1.3×10^13^ GC/ml) was used. To express iCre-P2A-mNeonGreen in pyramidal cells of *Nlgn2^f/f^;Sst^Flpo/+^;Rosa26^FSF-tdTomato/+^* mice at P0–2, 100 nl of AAV9-CaMKIIα-iCre-P2A-mNeonGreen (1.7×10^10^ GC/ml) was used. To densely express EGFP-Cre in neurons of *Nlgn2^f/f^* and *Nlgn2^+/+^* mice at P30, 200 nl of AAV9-hSyn-HI-EGFP-Cre (2.5×10^11^ GC/ml) was used.

### Double fluorescence *in situ* hybridization

A digoxigenin (DIG)-labeled RNA antisense probe against mouse *Nlgn2* and fluorescein (FITC)-labeled RNA antisense probes against *tdTomato* and mouse *Sst*, *Vip*, and *Pv* were generated by *in vitro* transcription using cDNA templates and RNA DIG-or FITC-labeling kits (Sigma, catalog # 11277073910 or 11685619910, respectively). The DNA templates were made by PCR amplification from a plasmid pCMV6-mNlgn2(A+)-mycDDK (Origene, catalog #MR222168, GenBank Accession NM_198862) for the *Nlgn2* probe, a plasmid pAAV-EF1α-DIO-tdTomato for the *tdTomato* probe, or mouse brain cDNA for the *Sst*, *Vip*, and *Pv* probes, with a T3 promoter (AATTAACCCTCACTAAAGGG) added at the 5’ end of the PCR forward primers and a T7 promoter (TAATACGACTCACTATAGGG) at the 5’ end of the PCR reverse primers. The *Nlgn2* probe was designed to span exons 4–6. The sequences of *tdTomato*, *Sst*, *Vip*, and *Pv* probes were from Allen Brain Atlas (http://mouse.brain-map.org). All probe sequences are listed in **Supplementary File 1**.

Double fluorescence *in situ* hybridization (DFISH) was performed by the RNA *In Situ* Hybridization Core at Baylor College of Medicine using an automated robotic platform and procedures as described previously {Yaylaoglu:2005bb} with minor modifications for double ISH. Briefly, fresh-frozen brains were embedded in optimal cutting temperature (OCT) compound and cryosectioned at 14-µm thickness. Two probes were hybridized to brain sections simultaneously (*Nlgn2*/*Sst*, *Nlgn2*/*Vip*, *Nlgn2*/*Pv*, or *Nlgn2*/*tdTomato*) in hybridization buffer (Ambion, catalog #B8807G). Sections were washed with standard saline citrate stringency solution (SSC; 0.15 M NaCl, 0.015 M sodium citrate) to remove unbound and non-specifically bound probes. To visualize the DIG-labeled probe, brain sections were incubated for 30 minutes with a horse radish peroxidase (HRP)-conjugated sheep anti-DIG primary antibody (Sigma, catalog #11207733910) diluted at 1/500 in Tris-NaCl blocking buffer (TNB; 100 mM Tris, 150 mM NaCl, 0.5% (w/v) blocking reagent (Perkin Elmer, catalog #FP1012), pH 7.6). After washes in Tris-NaCl-Tween (TNT; 10 mM Tris-HCl, pH 8.0, 150 mM NaCl and 0.05% TWEEN 20) buffer, brain sections were then developed with tyramide-Cy3 Plus (Akoya Biosciences, catalog #NEL744001KT, 1/50 dilution in amplification diluent, 15 minutes). After washes in TNT buffer, the remaining HRP activity was quenched by a 10-minute incubation in 0.2 M HCl. Sections were then washed in TNT, blocked in TNB for 15 minutes before incubation with an HRP-conjugated sheep anti-FITC antibody (Sigma, catalog #11426346910) diluted at 1/500 in TNB for 30 minutes. After washes in TNT, the FITC-labeled probe was visualized using tyramide-FITC Plus (Akoya Biosciences, catalog #NEL741001KT, 1/50 dilution in amplification diluent, 15 minutes). The slides were washed in TNT and stained with 4’,6-diamidino-2-phenylindole (DAPI; Invitrogen, catalog #D3571), washed again, removed from the machine, and mounted in ProLong Diamond (Invitrogen, catalog #P36961).

To compare *Nlgn2* mRNA levels in Sst, Vip, or Pv cells between 8-week-old *Nlgn2^-/-^* mice and sex matched *Nlgn2^+/+^* control mice, sagittal brain sections were cut from 2 to 3 mm from the lambda and stained in parallel for *Nlgn2/Sst*, *Nlgn2/Vip*, or *Nlgn2/Pv*, respectively. To compare *Nlgn2* mRNA levels in Sst cells between *Nlgn2^f/f^;Sst^Cre/+^*;*Rosa26^Ai14/+^*and *Nlgn2^+/+^;Sst^Cre/+^;Rosa26^Ai14/+^* mice or in Vip cells between *Nlgn2^f/f^;Vip^Cre/+^*;*Rosa26^Ai14/+^*and *Nlgn2^+/+^;Vip^Cre/+^;Rosa26^Ai14/+^* mice at P14 and P21, sagittal brain sections were cut from 2 to 3 mm from the lambda and stained in parallel for *Nlgn2*/*tdTomato.* To compare *Nlgn2* mRNA levels in Pv cells between *Nlgn2^f/f^;Pv^Cre/+^;Rosa26^Ai14/+^* and *Nlgn2^+/+^;Pv^Cre/+^;Rosa26^Ai14/+^* mice at P14, sagittal brain sections were cut from 1 to 2 mm from the lambda. To compare *Nlgn2* mRNA levels in Sst cells between *Nlgn2^f/f^;Sst^Flpo/+^*;*Rosa26^Ai14/+^*and *Nlgn2^+/+^;Sst^Flpo/+^;Rosa26^Ai14/+^* mice or in Vip cells between *Nlgn2^f/f^;Vip^Flpo/+^*;*Rosa26^Ai14/+^*and *Nlgn2^+/+^;Vip^Flpo/+^;Rosa26^Ai14/+^* mice that were injected with AAV expressing iCre (see above), coronal brain sections were cut at P14 from approximately −4 to −3 mm from the bregma to cover the location of AAV injection and stained in parallel for *Nlgn2*/*tdTomato*.

### DFISH fluorescence imaging and analysis

Fluorescence images of the visual cortex were acquired on an Sp8X Confocal Microscope (Leica) using a 40x oil objective. 3–4 brain sections per mouse were imaged based on the density of Sst, Vip, or Pv cells in the section. Approximately 40–60 images were acquired per tile scan with a 5% overlap between images for tiling. The z-stack (∼20 optical sections, 0.6-µm step) contained the entire thickness of the brain section.

For analysis, the images were opened in ImageJ and z-projected using the “Sum Slices” function and then converted to a .ims format for analysis in Imaris 9.7 (Oxford Instruments). To compare *Nlgn2* mRNA levels between *Nlgn2^+/+^* and *Nlgn2^−/−^* mice for the Sst, Vip, or Pv cells in layer 2/3, neurons were identified based on DAPI-positive nuclei using the “Surfaces” function with the following parameters: smooth with surface grain size of 0.455 µm, eliminate background with diameter of largest sphere = 1.70 µm, minimum threshold value of 300, and number of voxels between 1000 to 2500. Surfaces were then manually inspected to rule out glial nuclei (i.e., elongated nuclei) and exclude any surfaces that contained two nuclei. The mean Nlgn2 intensity was then calculated for each surface that was positive for *Sst*, *Vip*, or *Pv*. The background intensity in the *Nlgn2* channel was calculated by averaging the mean intensity in *Nlgn2* channel from 5 ovals of area ∼150 µm^2^ in the intercellular space of layer 2/3. This value was subtracted from the mean *Nlgn2* intensity of the *Sst*, *Vip*, or *Pv*-positive surfaces.

To analyze *Nlgn2* mRNA levels for *tdTomato*-positive Sst, Vip, or Pv cells following Cre-mediated deletion of *Nlgn2*, a 3 x 3 x 1 median filter was applied to the *tdTomato* channel of each image to decrease the background. The *tdTomato*-positive cells in layer 2/3 were identified using the “Surfaces” function with the following parameters: smooth with surface grain size of 1.00 µm, eliminate background with diameter of largest sphere = 5.00 µm, minimum threshold value of 50, and number of voxels about 1000. *tdTomato* surfaces were then manually inspected to ensure that 1) each surface represented a real cell as evidenced by colocalization of the *tdTomato* signal and a DAPI-positive nucleus and 2) each surface represented the *Nlgn2* mRNA from only a single cell. If the nucleus of a *tdTomato* surface had another nucleus located within 5 µm of it, then this *tdTomato* surface was excluded from the analysis due to the potential signal contamination. After confirming the accuracy of the *tdTomato* surfaces, the mean intensity of the *Nlgn2* channel was calculated for each surface. The background intensity in the *Nlgn2* channel was calculated by averaging the mean intensity in *Nlgn2* channel from 5 ovals of area ∼150 µm^2^ (roughly the same area as *tdTomato* surfaces) in the intercellular space of layer 2/3. This value was subtracted from the mean *Nlgn2* intensity of the *tdTomato* surfaces. Due to the variability in *Nlgn2* intensity between different staining batches and mice, surfaces from each section were then normalized to the mean *Nlgn2* intensity of all cells in layer 2/3, which served as an internal control for staining variability. This was done by using the “Surfaces” function to generate a surface from the DAPI channel containing all cells in layer 2/3 with the following parameters: smooth with surface grain size of 1.00 µm, eliminate background with diameter of largest sphere = 1.70 µm, minimum threshold of 5, and number of voxels between 1000 and 5000. Approximately 1000 surfaces were generated for each section. The mean *Nlgn2* intensity minus the background was calculated for each surface and averaged across all surfaces to determine the normalization value for each section. The mean *Nlgn2* intensity in each *tdTomato* surface was then divided by the normalization value to determine the final normalized mean *Nlgn2* intensity for each *tdTomato*-positive cell.

### Brain slice electrophysiology

Mice were anesthetized by an intraperitoneal injection of a ketamine and xylazine mix (80 mg/kg and 16 mg/kg, respectively) and transcardially perfused with cold (0–4°C) slice cutting solution containing 80 mM NaCl, 2.5 mM KCl, 1.3 mM NaH_2_PO_4_, 26 mM NaHCO_3_, 4 mM MgCl_2_, 0.5 mM CaCl_2_, 20 mM D-glucose, 75 mM sucrose, and 0.5 mM sodium ascorbate (315 mosmol, pH 7.4, saturated with 95% O_2_/5% CO_2_). Brains were removed and sectioned in the cutting solution with a VT1200S vibratome (Leica) to obtain 300 mm coronal slices. Slices containing the primary visual cortex were collected and incubated in a custom-made interface holding chamber saturated with 95% O_2_/5% CO_2_ at 34°C for 30 min and then at room temperature for 20 min to 6 hr until they were transferred to the recording chamber.

Recordings were performed on submerged slices in artificial cerebrospinal fluid (ACSF) containing 119 mM NaCl, 2.5 mM KCl, 1.3 mM NaH_2_PO_4_, 26 mM NaHCO_3_, 1.3 mM MgCl_2_, 2.5 mM CaCl_2_, 20 mM D-glucose, and 0.5 mM sodium ascorbate (305 mosmol, pH 7.4, saturated with 95% O_2_/5% CO_2_, perfused at 3 ml/min) at 32°C. For whole-cell recordings, a K^+^-based pipette solution containing 142 mM K^+^-gluconate, 10 mM HEPES, 1 mM EGTA, 2.5 mM MgCl^2^, 4 mM ATP-Mg, 0.3 mM GTP-Na, 10 mM Na_2_-phosphocreatine (295 mosmol, pH 7.35) or a Cs^+^-based pipette solution containing 121 mM Cs^+^-methanesulfonate, 10 mM HEPES, 10 mM EGTA, 1.5 mM MgCl_2_, 4 mM ATP-Mg, 0.3 mM GTP-Na, 10 mM Na_2_-phosphocreatine, and 2 mM QX314-Cl (295 mosmol, pH 7.35) was used. Membrane potentials were not corrected for liquid junction potential (experimentally measured as 12.5 mV for the K^+^-based pipette solution and 9.5 mV for the Cs^+^-based pipette solution).

Neurons were visualized with video-assisted infrared differential interference contrast imaging and fluorescent neurons were identified by epifluorescence imaging under a water immersion objective (40x, 0.8 numerical aperture) on an upright SliceScope Pro 1000 microscope (Scientifica) with an infrared IR-1000 CCD camera (DAGE-MTI). Data were acquired at 10 kHz and low-pass filtered at 4 kHz with an Axon Multiclamp 700B amplifier and an Axon Digidata 1550 or 1440 Data Acquisition System under the control of Clampex 10.7 (Molecular Devices). For the photostimulation of ChR2-expressing neurons, blue light (455 or 470 nm) was emitted from a collimated light-emitting diode (LED). The LED was driven by a LED driver (Mightex or Thorlabs) under the control of an Axon Digidata 1550 or 1440 Data Acquisition system and Clampex 10.7. Light was delivered through the reflected light fluorescence illuminator port and the 40x objective. Data were analyzed offline using AxoGraph X.

To record spontaneous postsynaptic currents (sPSCs), pyramidal cells or interneurons were recorded under voltage-clamp mode using the Cs^+^-based pipette solution at the reversal potential of inhibition (−70 mV) for sEPSCs or the reversal potential of excitation (+10 mV) for sIPSCs. Cells were recorded for 5–10 minutes for sEPSCs and sIPSCs. To detect sPSCs, data were digitally low-pass filtered at 2 kHz offline and events were detected by a scaled-template algorithm in AxoGraph X {Clements:1997he}. Different templates were used for detecting sPSCs in different cell types. For pyramidal cells, the parameters for sEPSCs are: length, 3 ms; baseline, 10 ms; amplitude, −2 pA; rise time, 0.6 ms; decay time, 3 ms; threshold, 3xSD, and the parameters for sIPSCs are: length, 3 ms; baseline, 20 ms; amplitude, 2 pA; rise time, 0.6 ms; decay time, 10 ms; threshold, 2.5xSD. For Sst cells, the parameters for sEPSCs are: length, 3 ms; baseline, 10 ms; amplitude, −2 pA; rise time, 0.4 ms; decay time, 1.9 ms; threshold, 3.25xSD, and the parameters for sIPSCs are: length, 3 ms; baseline, 20 ms; amplitude, 2 pA; rise time, 0.6 ms; decay time, 13 ms; threshold, 3.3xSD. For Vip cells, the parameters for sEPSCs are: length, 3 ms; baseline, 10 ms; amplitude, −2 pA; rise time, 0.43 ms; decay time, 2.95 ms; threshold, 3xSD, and the parameters for sIPSCs are: length, 3 ms; baseline, 15 ms; amplitude, 2 pA; rise time, 0.34 ms; decay time, 7.5 ms; threshold, 3xSD. For Pv cells, the parameters for sEPSCs are: length, 1.5 ms; baseline, 5 ms; amplitude, −2 pA; rise time, 0.2 ms; decay time, 1 ms; threshold, 3.25xSD, and the parameters for sIPSCs are: length, 3 ms; baseline, 10 ms; amplitude, 2 pA; rise time, 0.27 ms; decay time, 3.7 ms; threshold, 3xSD.

To record photostimulation-evoked inhibitory postsynaptic currents (eIPSCs), a pair of control and iCre-positive pyramidal cells or Sst cells that were within were within 50 µm of each other were recorded under voltage-clamp mode using the Cs^+^-based pipette solution at the reversal potential of excitation (+10 mV). Blue light pulse duration (0.5–5 ms) and intensity (1.1–5.5 mW/mm^2^) were adjusted for each recording to evoke small (to minimize voltage-clamp errors) but reliable monosynaptic IPSCs. Light pulses were delivered at 30-s inter-trial intervals. eIPSC amplitudes were measured from the average of 5–7 trials.

To record unitary connections between Sst cells and pyramidal cells, a control and an iCre-positive pyramidal cell were recorded under voltage-clamp mode at the reversal potential of excitation (+10 mV) using the Cs^+^-based pipette solution, and a nearby (within 50 µm of both pyramidal cells) Sst cell was identified by the Flpo-dependent expression of tdTomato and recorded under current-clamp mode using the K^+^-based pipette solution. Action potentials were elicited in Sst cells by a train of 6 depolarizing current steps (2 ms, 1–2 nA) at 10 Hz with 15-s inter-trial intervals. Unitary IPSC (uIPSC) amplitudes were measured from the first IPSCs of the average of 50 trials. A Sst cell was considered to be connected with a pyramidal cell if the average uIPSC amplitude was at least three times of the baseline standard deviation.

### Fluorescence imaging of live brain slices and analysis

Fluorescence images were acquired from 300 µm-thick live brain slices immediately following electrophysiological recordings. Images were acquired on an Axio Zoom.V16 Fluorescence Stereo Zoom Microscope (Zeiss) at 20x (whole brain) or 63x (visual cortex) to visualize the fluorescent cells. Images were manually analyzed in ImageJ to quantify the numbers of tdTomato-positive cells and GFP-positive cells.

### Neural network modeling

#### Firing-rate-based population model

A simple recurrent network consisting of one population of excitatory cells (E cells) and one population of inhibitory cells (I cells) was constructed. The state of the network is characterized by *r_E_*(*t*), *r_I_*(*t*), which are the population firing rates of E cells and I cells, respectively. The temporal evolution of *r_E_*(*t*), *r_I_*(*t*) is given by temporally coarse-grained equations {Wilson:1972kj}:

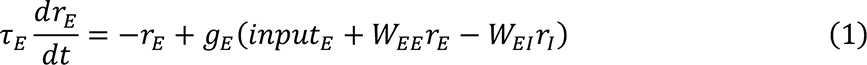

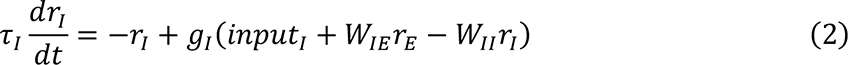

Here *input_E_*, *input_I_* are the tonic external inputs to each cell in the E population and I population, respectively. *τ_E_*, *τ_I_* are the time constants. *W_EI_* is the strength of connection from I population to E population (*I* → *E*). Similarly, *W_EE_*, *W_IE_*, and *W_II_* represent *E* → *E*, *E* → *I*, and *I* → *I* connections, respectively. *W_EE_*, *W_EI_*, *W_IE_*, and *W_II_* > 0. The response functions of each population, *g_E_*(*x*) and *g_I_*(*x*), are given by:

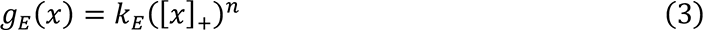

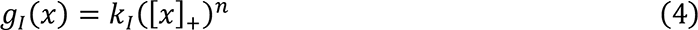

Here *k_E_* and *k_I_* are positive constants. [. ]_+_ denotes the rectifier: [*x*]_+_ = *x* if *x* > 0; = 0 otherwise. *n*, the power of the response function, is taken identical for E and I populations for simplicity. Note that here we take *n* > 1 to model a stabilized supralinear network (SSN), which has been proposed to explain various nonlinear cortical computation {Ahmadian:2013kq}.

#### Linearized equations about the fixed points

In the small neighborhood of the fixed points 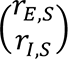 satisfying 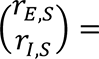 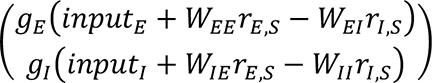, with external input fixed, the model can be linearized to:

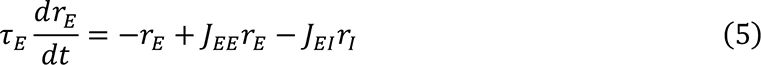

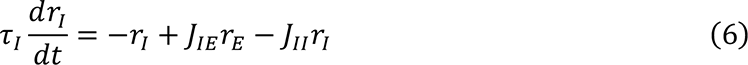

Where

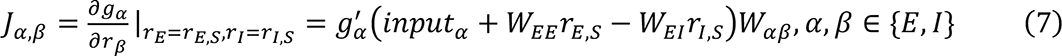

Note that *r_E_*(*t*) and *r_I_*(*t*) are now defined as the deviations from the fixed points (*r_E,S_*, *r_I,S_*). Equations (5) and (6) can be expressed in matrix forms:

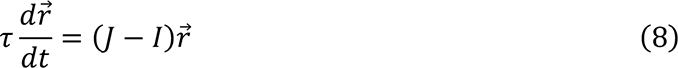

The vector of population firing rates 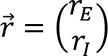, the connectivity matrix 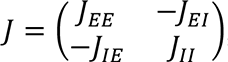, the time constant matrix 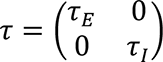, and the identity matrix 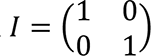.

#### Conditions for the network to operate as an inhibition-stabilized network (ISN)

A network would operate as an ISN if the following two conditions are met: (1) The network would be unstable without feedback inhibition, and (2) the feedback inhibition is sufficiently strong to stabilize the network. Condition (1) requires that the linearized network without feedback inhibition about the fixed points is unstable, which is equivalent to that 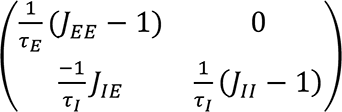 has at least one positive eigenvalue. The two eigenvalues of the matrix are 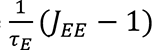 and 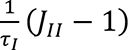. As 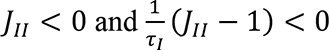, the instability of the network is equivalent to J*_EE_* > 1. Similarly, condition (2) requires that the linearized network with feedback inhibition about the fixed points is stable, which is equivalent to that 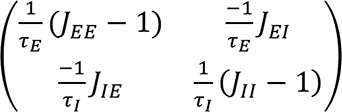 has two eigenvalues with negative real parts. This is in turn equivalent to that the matrix has a positive determinant and a negative trace, which reduces to 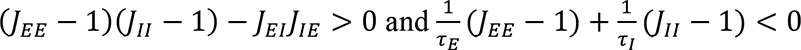.

#### Response of the network to changes in external inputs

To characterize the response of the network, how its stationary rate will change for a small change in external inputs was considered. For the stationary state of the network 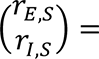 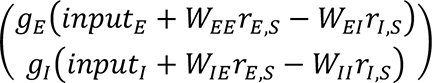, with a small perturbation 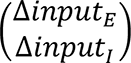 applied to the original external inputs 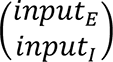, the network will reach a new stationary state 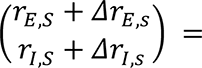 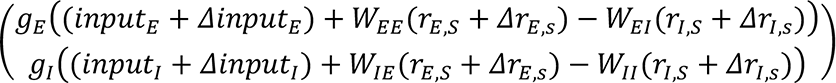. Under linearization,

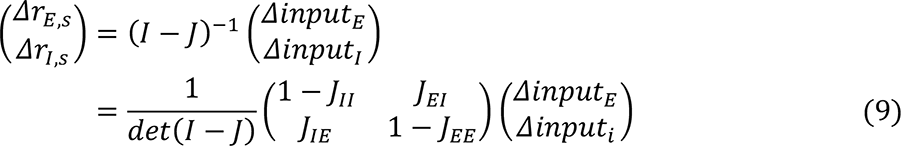

The paradoxical effect refers to that the modulation of the external input to I cells leads to a paradoxical change of their firing rate, i.e., Δ*input_I_* always has a negative effect on *Δr_I_*_,S_. In the regime of ISN, *det*(*I* − *J*) > 0. Therefore, the paradoxical effect is present only when 1 – *J_EE_* < 0. Note that although both exhibiting a paradoxical effect and the instability of the network in the absence of feedback inhibition require ‘1 – *J_EE_* < 0’, they refer to different stationary rates. Due to the nonlinearity of the response functions, these two properties are not equivalent.

#### Simulation method

Stationary states were determined by simulations using forward Euler method with *dt* = 0.1. Multiple initial conditions were applied to confirm the independence of final stationary state from initial condition.

### Experimental study design and statistics

Estimation of the sample size was made based on the previous studies {Xue:2014dl, Chen:2020ga} that used similar assays and pilot experiments. They are within the range that is generally accepted in the field. All experiments were performed and analyzed blind to the genotypes. Both male and female mice were included in experiments. No data point was excluded.

For DFISH and electrophysiology experiments, all reported sample numbers (*n*) represent the number of total cells followed by the number of mice. For health monitoring, all reported sample numbers represent tested mice. Statistical analyses were performed with Prism 9 (Graphpad Software). Anderson-Darling test, D’Agostino-Pearson, Shapiro-Wilk, and Kolmogorov-Smirnov tests were used to determine if data were normally distributed. If all data within one experiment passed all four normality tests, then the statistical test that assumes a Gaussian distribution was used. Otherwise, the statistical test that assumes a non-Gaussian distribution was used. The details of all statistical tests, numbers of replicates, and *P* values are reported in **Supplementary File 2**.

## Materials and data availability

All new plasmids generated in this study will be available from Addgene (see DNA constructs section). All data generated and analyzed in this study are included in the manuscript.

---

**Supplemental Figure 1 (Related to Figure 1):**
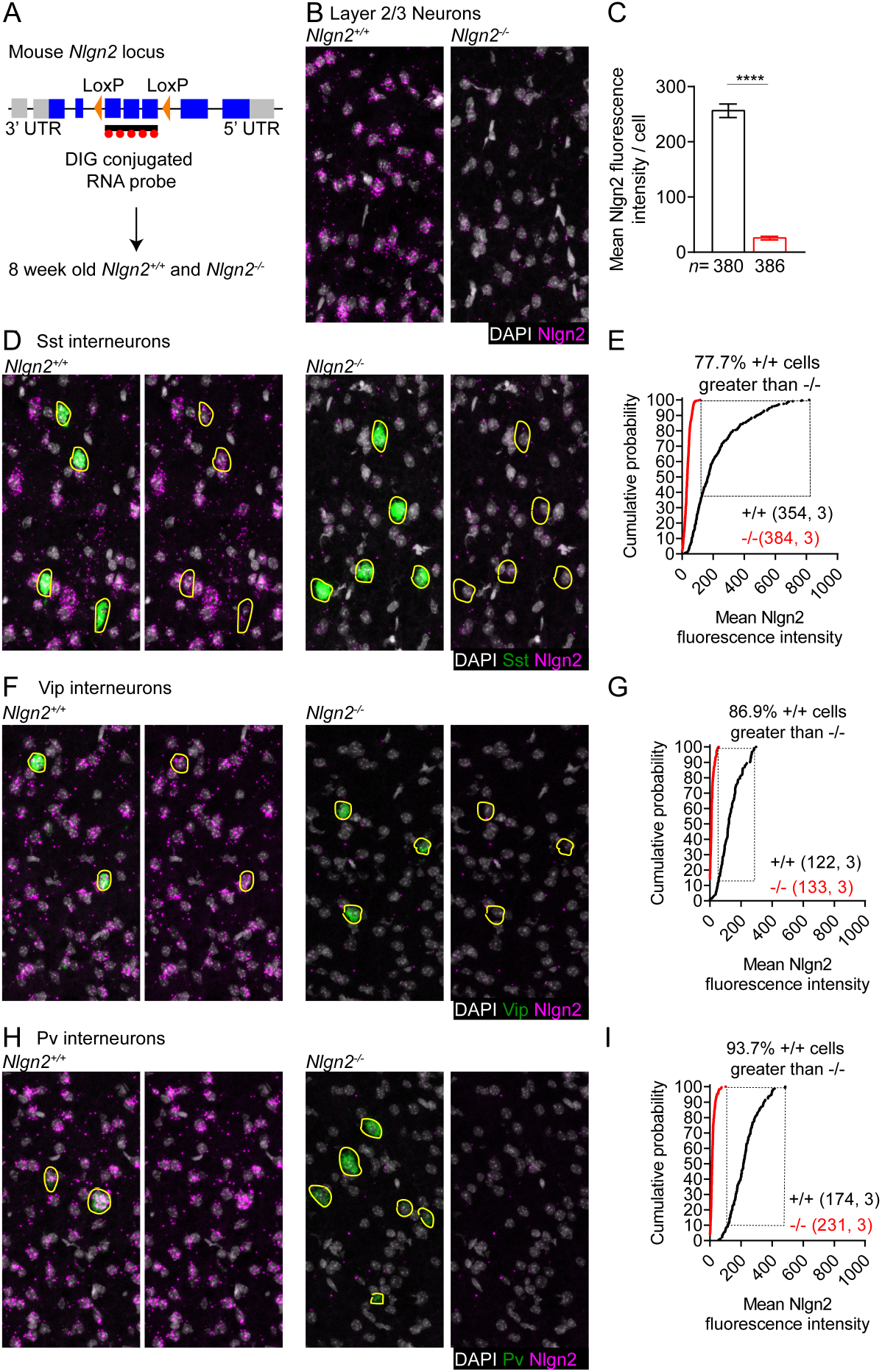
*Nlgn2* mRNA is expressed in Sst, Vip, and Pv interneurons (**A**) Genomic structure of the *Nlgn2* locus in mouse with LoxP sites flanking the region that is deleted following Cre mediated recombination. The area recognized by the *Nlgn2* ISH probe is also indicated. (**B**) Representative images of DFISH showing *Nlgn2* (magenta) and DAPI (gray) from 8-week-old *Nlgn2^+/+^* (left) and *Nlgn2^-/-^* (right) mice. (**C**) Summary data of the *Nlgn2* levels of layer 2/3 neurons. (**D**) Representative images of DFISH showing Nlgn2 (magenta), *Sst* (green), and DAPI (gray) in the visual cortex of 8-week-old *Nlgn2^+/+^*(left) and *Nlgn2^-/-^* (right) mice. Yellow outlines represent Sst positive cells. (**E**) Cumulative frequencies of Sst positive cells as a function of their *Nlgn2* levels. Dashed lines indicate that 77.7% of Sst positive cells from *Nlgn2^+/+^* mice are greater than all Sst positive cells in *Nlgn2^-/-^* mice. (**F,G**) As in (C,D), but for Vip positive cells. (**H,**I) As in (D,E), but for Pv positive cells. *N* represents # of cells followed by # of animals. Bar graph is mean ± S.E.M. ****, p<0.0001. Scale bar is 50 µm.

**Supplemental Figure 2 (Related to Figure 1):**
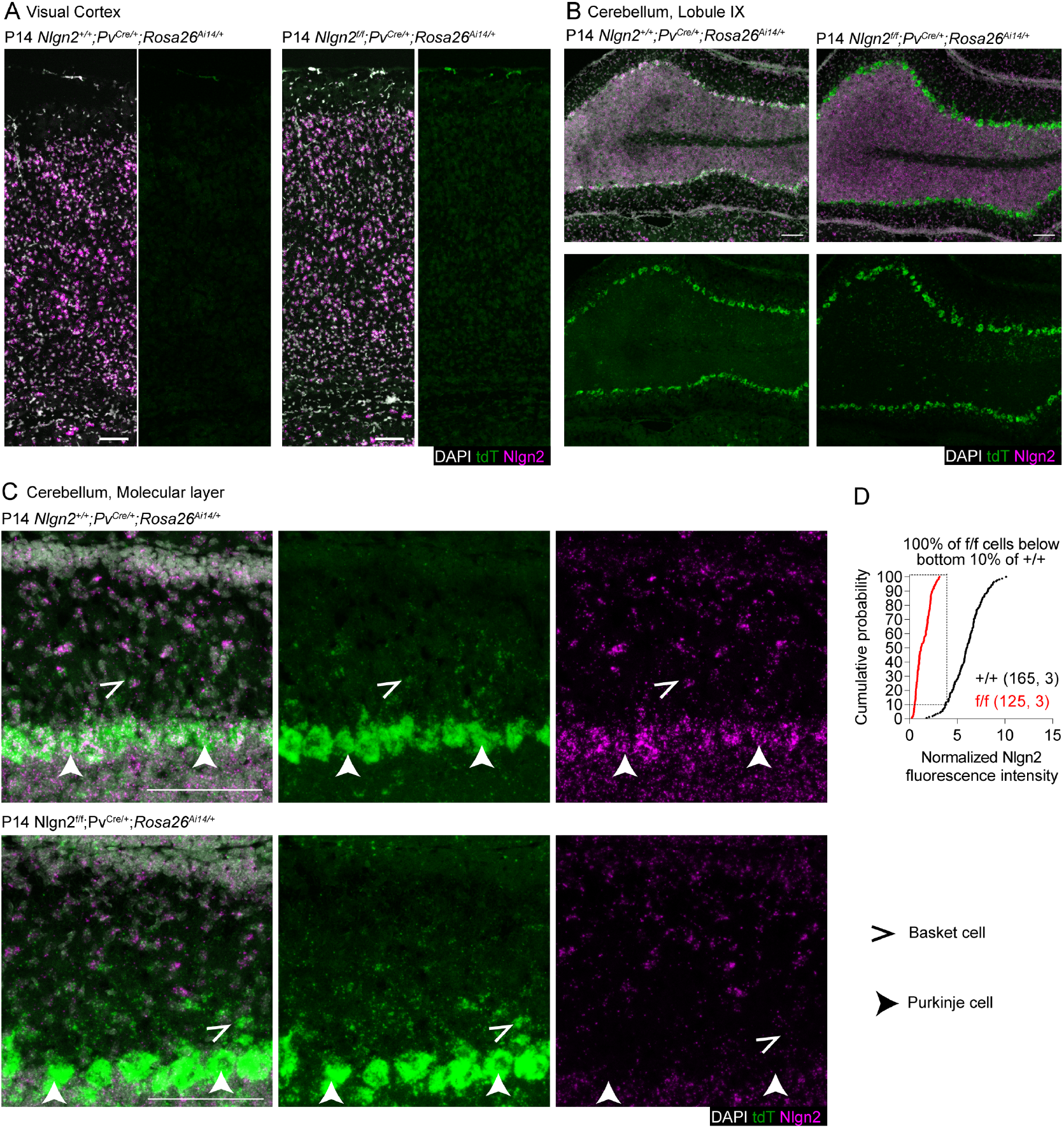
At P14 Pv-Cre is not expressed in visual cortex but efficiently deletes *Nlgn2* in cerebellum (**A**) Representative images of DFISH showing *Nlgn2* (magenta), *tdTomato* (green), and DAPI (gray) in the visual cortex of P14 *Nlgn2^+/+^;Pv^Cre/+^;Rosa26^Ai14/+^*(left) and *Nlgn2^f/f^*;*Pv^Cre/+^;Rosa26^Ai14/+^*(right) mice. (**B**) As in (A) but for lobule IX of the cerebellum. (**C**) As in (A), but for the molecular layer of lobule IX of the cerebellum. High magnification images of the molecular layer of lobule IX of the cerebellum. Closed arrowhead indicates Purkinje cells and open arrowhead indicates basket cells. (**D**) Cumulative frequencies of tdTomato positive Purkinje cells as a function of their *Nlgn2* levels. Dashed line indicates that 100% of tdTomato positive cells in *Nlgn2^f/f^*;*Pv^Cre/+^;Rosa26^Ai14/+^* mice are within the bottom 10% of tdTomato positive purkinje cells from *Nlgn2^+/+^;Pv^Cre/+^;Rosa26^Ai14/+^* mice. *N* represents # of cells followed by # of animals. Scale bar is 100 µm.

**Supplemental Figure 3 (Related to Figure 2):**
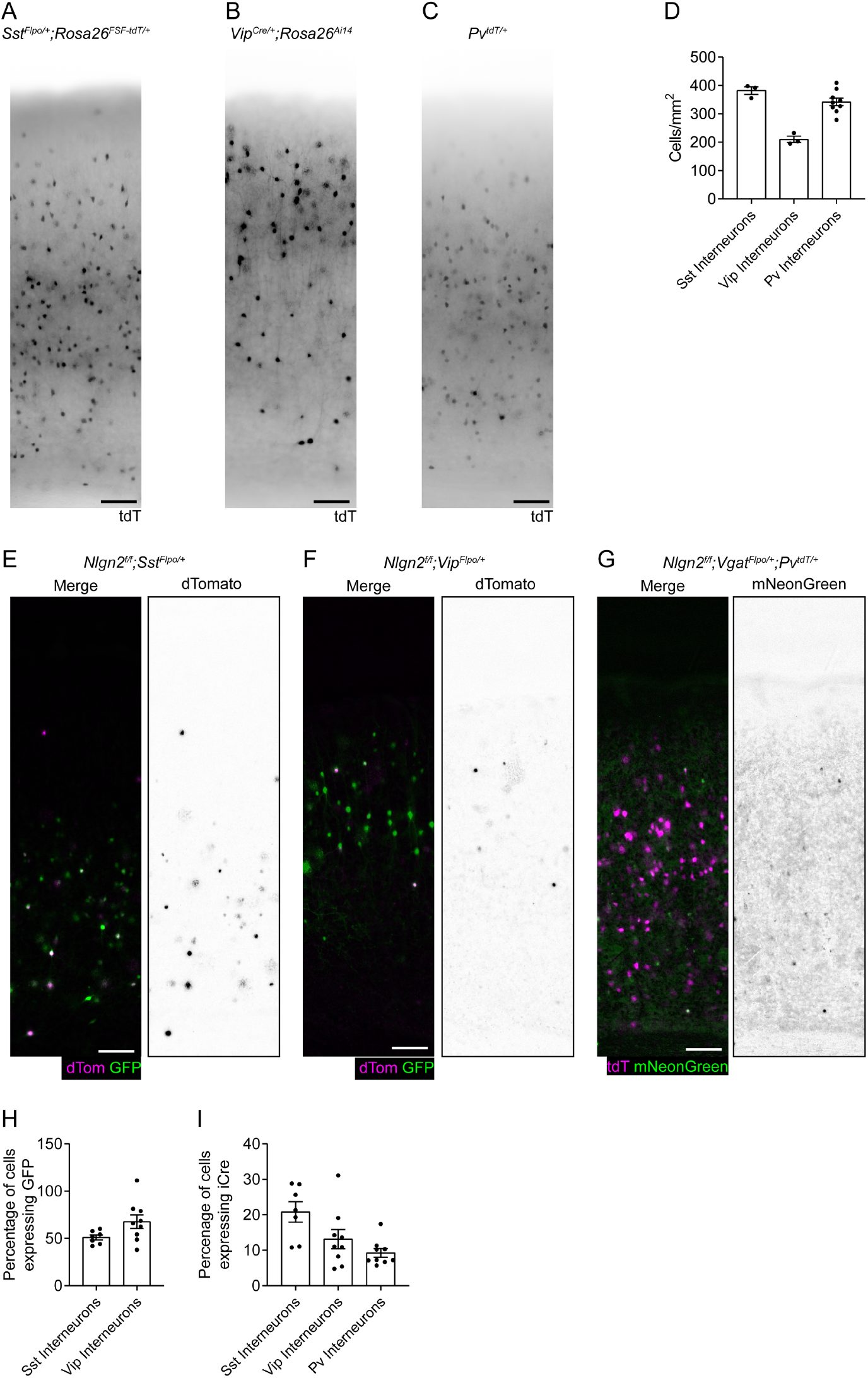
Determination of viral titer needed to infect 10% of each interneuron subtype with iCre. (**A**) Representative image of a 300 µm live section from visual cortex of a *Sst^Flpo/+^;Rosa26^FSF-tdT/+^*animal showing tdTomato signal in Sst interneurons. (**B**) As in (A), but for Vip interneurons in a *Vip^Cre/+^;Rosa26^Ai14/+^* animal. (**C**) As in (A), but for Pv interneurons in a *Pv^tdT/+^* animal. (**D**) Summary data of the number of tdT cells/mm^2^ for Sst interneurons in *Sst^Flpo/+^;Rosa26^FSF-tdT/+^* animals, Vip interneurons in *Vip^Cre/+^;Rosa26^Ai14/+^* animals, and Pv interneurons in *Pv^tdT/+^* animals. (**E**) Representative image of a 300 µm live section from visual cortex of a *Nlgn2^f/f^;Sst^Flpo/+^*animal injected at P0–P2 with AAV9-FRT-FLEX-iCre-P2A-3xNLS-dTomato and AAV9-FRT-FLEX-GFP showing GFP (green) and dTomato (magenta) merged (left) and dTomato alone (right). (**F**) As in (E), but for a *Nlgn2^f/f^;Vip^Flpo/^*^+^ animal. (**G**) Representative image of a 300 µm live section from visual cortex of a *Nlgn2f/f;VgatFlpo/+;PvtdT/+* animal injected at P0–P2 with AAV9-FRT-FLEX-iCre-P2A-3xNLS-mNeonGreen. (**H**) Summary data of the percentage of Sst or Vip interneurons expressing GFP following viral injection. (**I**) Summary data of the percentage of Sst, Vip, or Pv interneurons expressing iCre following viral injection. Bar graph is mean ± S.E.M. Each dot represents the average number from 2-5 sections for one animal. Scale bar is 100 µm.

**Supplemental Figure 4 (Related to Figure 2):**
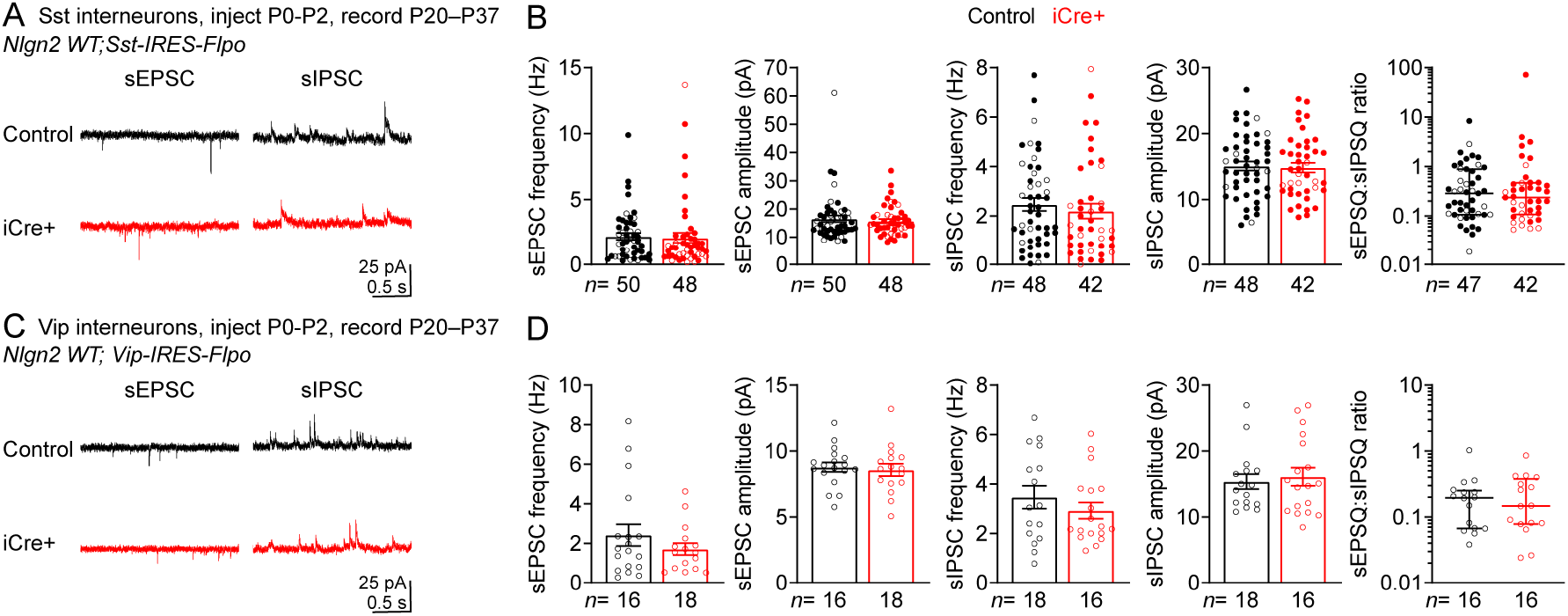
Flpo dependent Cre virus has no effect on inhibition onto Sst and VIP cells in *Nlgn2^+/+^* mice. (**A**) Representative trace of sEPSCs (left) and sIPSCs (right) from control (top) and iCre+ (bottom) Sst cells from *Nlgn2^+/+^;Sst^Flpo/+^* mice injected with AAV9-FRT-FLEX-iCre-P2A-3xNLS-dTomato and AAV9-FRT-FLEX-GFP at P0– P2. (**B**) Summary data of frequency and amplitude of sEPSCs, sIPSCs, and sEPSQ:sIPSQ ratio between control and iCre+ cells in *Nlgn2^+/+^;Sst^Flpo/+^* mice injected at P0–P2. (**C,D**) As in (A,B), but for VIP cells from *Nlgn2^+/+^;Vip^Flpo/+^* mice. Each filled (male) or open (female) circle represents one neuron. Bar graphs are mean ± S.E.M. except for sEPSQ:sIPSQ ratio which is median ± interquartile range.

**Supplemental Figure 5 (Related to Figure 2):**
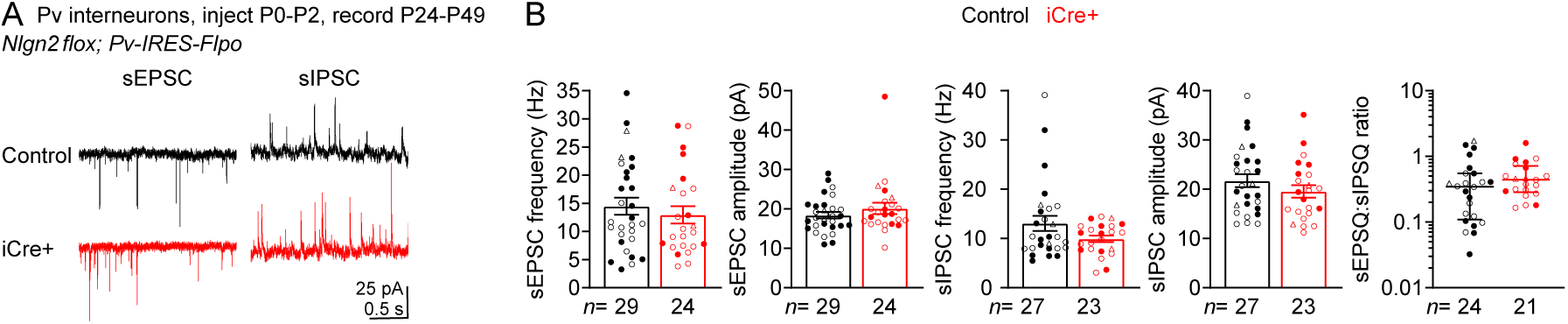
Targeting Pv cells using *Pv-IRES-Flpo* and Flpo dependent Cre virus has no effect on inhibition onto Pv cells. (**A**) Representative trace of sEPSCs (left) and sIPSCs (right) from control (top) and iCre+ (bottom) Pv cells from *Nlgn2^f/f^;Pv^Flpo/+^*mice injected with AAV9-FRT-FLEX-iCre-P2A-3xNLS-dTomato and AAV9-FRT-FLEX-GFP at P0–P2. (**B**) Summary data of frequency and amplitude of sEPSCs, sIPSCs, and sEPSQ:sIPSQ ratio between control and iCre+ cells in *Nlgn2^f/f^;Pv^Flpo/+^*mice injected at P0–P2. Each filled (male) or open (female) circle represents one neuron. Bar graphs are mean ± S.E.M. except for sEPSQ:sIPSQ ratio which is median ± interquartile range.

**Supplemental Figure 6 (Related to Figure 3):**
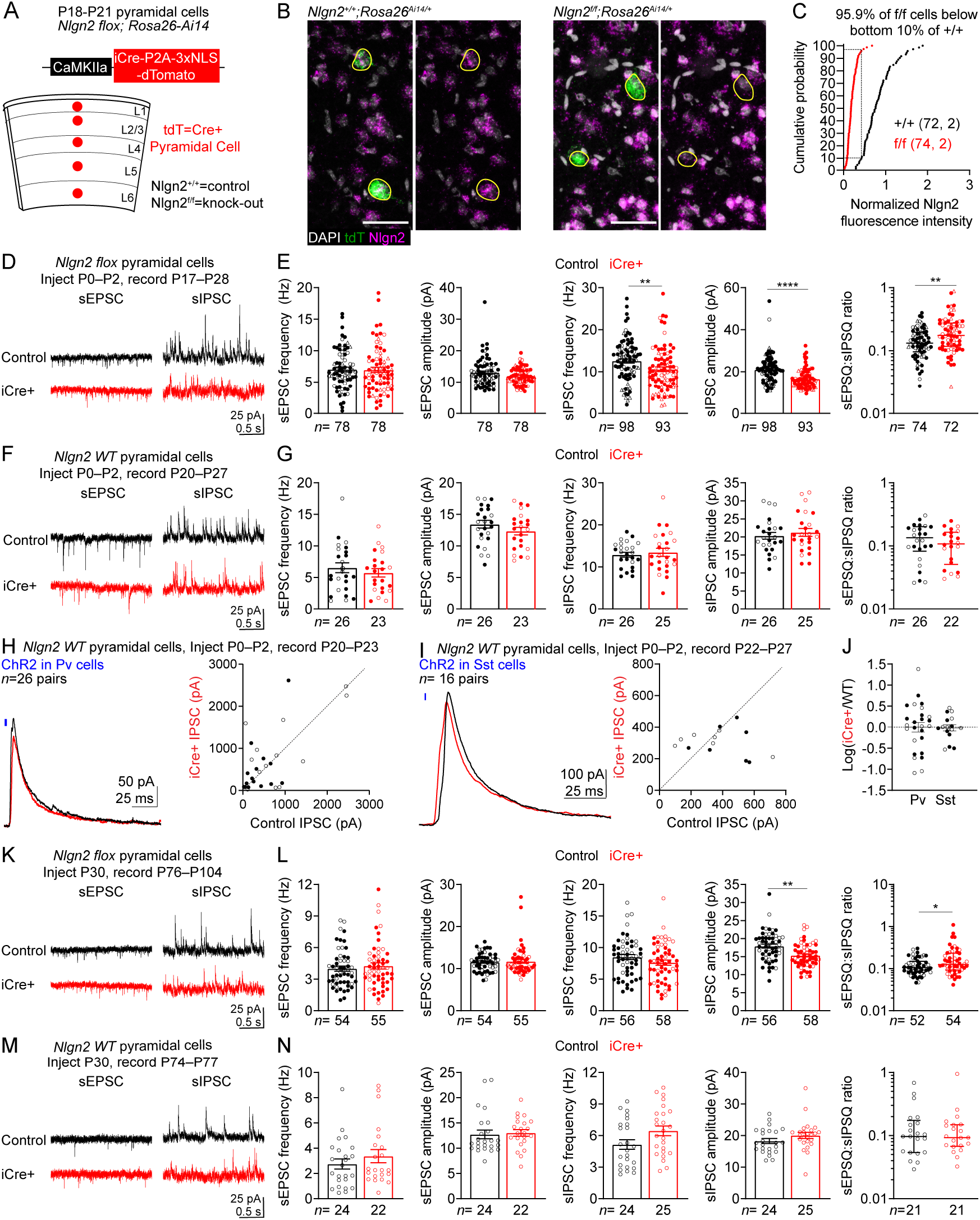
Early deletion of Nlgn2 in pyramidal cells disrupts inhibition but adult deletion has no affect. (**A**) Schematic illustrating genotypes, viral vectors, and experimental design for DFISH of *Nlgn2* in pyramidal cells. (**B**) Representative images of DFISH showing *Nlgn2* (magenta), *tdTomato* (green), and DAPI (gray) from P18–P21 *Nlgn2^f/f^;Rosa26^Ai14/+^* and *Nlgn2^+/+^;Rosa26^Ai14/+^*mice injected at mice injected at P0–P2 with CaMKIIa-iCre-P2A-3xNLS-dTomato. Yellow outlines represent tdTomato positive cells. (**C**) Cumulative frequencies of tdTomato positive cells as a function of their *Nlgn2* levels. Dashed lines indicate that 95.9% of tdTomato positive cells from *Nlgn2^f/f^;Rosa26^Ai14/+^* mice are within the bottom 10% of tdTomato positive cells from *Nlgn2^+/+^;Rosa26^Ai14/+^* mice. (**D**) Representative trace of sEPSCs (left) and sIPSCs (right) from control (top) and iCre+ (bottom) layer 2/3 pyramidal cells from *Nlgn2^f/f^*mice injected at P0–P2 with AAV9-CaMKIIa-iCre-P2A-3xNLS-dTomato. (**E**) Summary data of frequency and amplitude of sEPSCs, sIPSCs, and sEPSQ:sIPSQ ratio between control and iCre+ layer 2/3 pyramidal cells in *Nlgn2^f/f^* mice injected at P0–P2. (**F,G**) As in (D,E), but for *Nlgn2^+/+^* mice. (**H**) (Left) Representative trace of a Pv evoked IPSC onto a control (black) and iCre+ (red) layer 2/3 pyramidal cell in *Nlgn2^+/+^;Pv^Flpo/+^* mice injected with AAV9-CaMKIIa-iCre-P2A-3xNLS-dTomato and AAV9-FRT-FLEX-ChR2. (Right) Summary data of the amplitude of Pv evoked IPSCs onto pairs of control and iCre+ layer 2/3 pyramidal cells from *Nlgn2^+/+^;Pv^Flpo/+^*mice following P0–P2 injection. (**I**) As in (H) but for Sst evoked IPSCs from *Nlgn2^+/+^;Sst^Flpo/+^* mice. (**J**) Summary data of the log ratio of the amplitude of Pv and Sst evoked IPSCs onto pairs of control and iCre+ layer 2/3 pyramidal cells. (**K,L**) As in (D.E) but following P30 injection of CaMKIIa-iCre-P2A-3xNLS-dTomato. (**M,N**) As in (K,L), but for *Nlgn2^+/+^* mice. N number represents # of cells followed by number of animals for cumulative frequency graphs or # of cells for physiology summary data. Each filled (male) or open (female) circle represents one neuron. Open triangle indicates sex unknown. Bar graphs are mean ± S.E.M. except for sEPSQ:sIPSQ ratio which is median ± interquartile range. *, p<0.05; **, p<0.005; ****, p<0.0001. Scale bar is 50 µm.

**Supplemental Figure 7 (Related to Figure 3):**
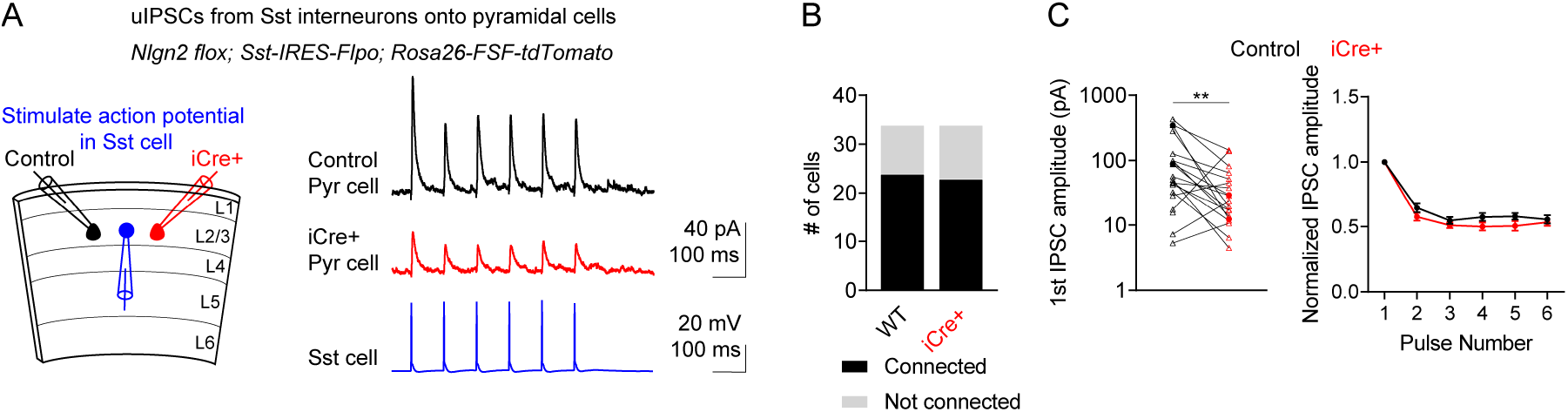
Unitary IPSC between Sst cells and layer 2/3 pyramidal cells are reduced following deletion of *Nlgn2*. (**A**) (Left) Schematic illustrating experimental design for recording unitary IPSC from Sst cell onto pair of control and iCre+ layer 2/3 pyramidal cells. (Right) Representative trace of Sst evoked IPSC onto control (black) and iCre+ (red) layer 2/3 pyramidal cells from *Nlgn2^f/f^;Sst^Flpo/+^;Rosa26^FSF-TdT/+^*mice following P0– P2 injection of AAV9-CaMKIIa-iCre-P2A-3xNLS-mNeonGreen. Blue trace indicates action potentials evoked by stimulating Sst cell. (**B**) Summary data of the rate of connectivity between control and iCre+ pyramidal cells and nearby Sst cells. (**C**) (Left) Summary data of the amplitude of the first IPSC evoked by stimulating Sst cell between control and iCre+ pyramidal cells. Lines connect pairs of control and iCre+ cells. (Right) Summary data of the amplitude of the IPSC evoked by each stimulation of the Sst cell normalized to the amplitude of the first IPSC evoked for a given cell between control and iCre+ pyramidal cells. Each filled (male) or open (female) circle represents one neuron. Open triangle indicates sex unknown. XY plot is mean ± S.E.M. **, p<0.005.

**Supplemental Figure 8 (Related to Figure 3):**
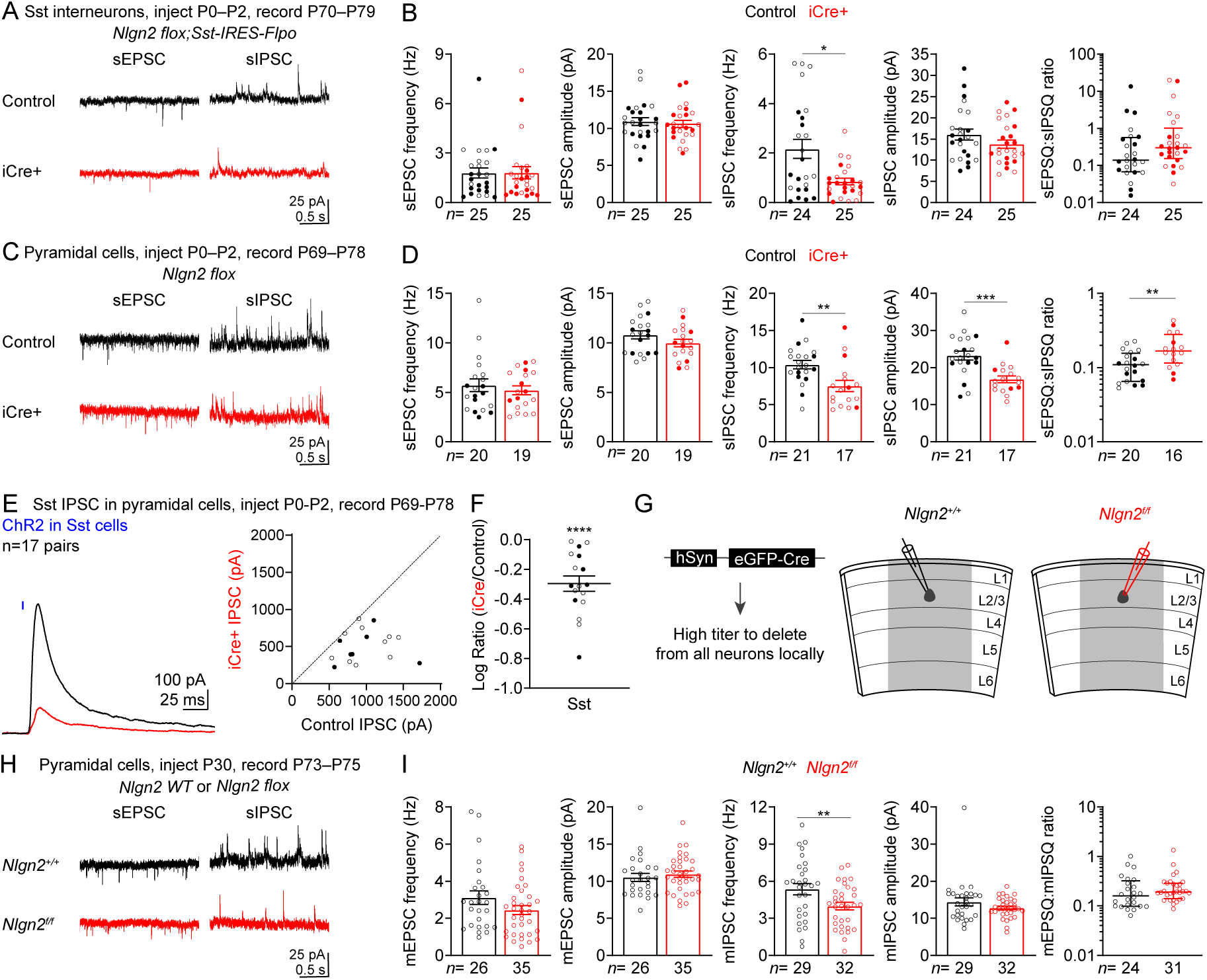
Recording age does not impact *Nlgn2* mediated decrease in inhibition but density of *Nlgn2* knock-out cells does. (**A**) Representative traces of sEPSCs (left) and sIPSCs (right) from control and iCre+ Sst cells from *Nlgn2^f/f^;Sst^Flpo/+^* mice injected at P0–P2 with AAV9-FRT-FLEX-iCre-P2A-3xNLS-dTomato and AAV9-FRT-FLEX-GFP and recorded at ∼10 weeks of age. (**B**) Summary data of frequency and amplitude of sEPSCs, sIPSCs, and sEPSQ:sIPSQ ratio between control and iCre+ Sst cells from *Nlgn2^f/f^;Sst^Flpo/+^* mice injected at P0–P2 and recorded at ∼10 weeks of age. (**C,D**) As in (A,B), but for layer 2/3 pyramidal cells from *Nlgn2^f/f^*;*Sst^Flpo/+^*mice injected at P0–P2 with AAV9-CaMKIIa-iCre-P2A-3xNLS-dTomato and FFLEX-ChR2 and recorded at ∼10 weeks of age. (**E**) (Left) Representative trace of an Sst evoked IPSC onto control (black) and iCre+ (red) layer 2/3 pyramidal cells from *Nlgn2^f/f^*;*Sst^Flpo/+^*mice injected at P0–P2 with AAV9-CaMKIIa-iCre-P2A-3xNLS-dTomato and AAV9-FRT-FLEX-ChR2. (Right) Summary data of the amplitude of Sst evoked IPSCs onto pairs of control and iCre+ layer 2/3 pyramidal cells from *Nlgn2^f/f^*;*Sst^Flpo/+^*mice injected at P0–P2 and recorded at ∼10 weeks of age.(**F**) Summary data of the log ratio of the amplitude of Sst evoked IPSCs from pairs of control and iCre+ layer 2/3 pyramidal cells. (**G**) Schematic illustrating the viral vectors, genotypes, and experimental design for dense deletion of *Nlgn2* from all cells within local region of visual cortex. (**H**) Representative traces of sEPSCs (left) and sIPSCs (right) from layer 2/3 pyramidal cells of *Nlgn2^+/+^*(black) or *Nlgn2^f/f^* (red) mice following P30 injection of hSyn-eGFP-Cre. (**I**) Summary data of frequency and amplitude of sEPSCs, sIPSCs, and sEPSQ:sIPSQ ratio between layer 2/3 pyramidal cells of *Nlgn2^+/+^*and *Nlgn2^f/f^* mice. Each filled (male) or open (female) circle represents one neuron. Bar graphs are mean ± S.E.M. except for sEPSQ:sIPSQ ratio which is median ± interquartile range. *, p<0.05; **, p<0.005; ***, p<0.0005; ****, p<0.0001.

## References

Adesnik, H. and Naka, A. (2018) ‘Cracking the Function of Layers in the Sensory Cortex’, Neuron. Elsevier Inc., 100, pp. 1028–1043. doi: 10.1016/j.neuron.2018.10.032.

Ahmadian, Y., Rubin, D. B. and Miller, K. D. (2013) ‘Analysis of the Stabilized Supralinear Network’, Neural Computation, 25, pp. 1994–2037. doi: 10.1162/NECO.

Almási, Z. et al. (2019) ‘Distribution patterns of three molecularly defined classes of gabaergic neurons across columnar compartments in mouse barrel cortex’, Frontiers in Neuroanatomy, 13(April), pp. 1–12. doi: 10.3389/fnana.2019.00045.

Amit, D. J. and Brunel, N. (1997) ‘Model of global spontaneous activity and local structured activity during delay periods in the cerebral cortex’, Cerebral Cortex, 7(3), pp. 237–252. doi: 10.1093/cercor/7.3.237.

Antoine, M. W. et al. (2019) ‘Increased Excitation-Inhibition Ratio Stabilizes Synapse and Circuit Excitability in Four Autism Mouse Models’, Neuron. Elsevier Inc., 101(4), pp. 648–661.e4. doi: 10.1016/j.neuron.2018.12.026.

Ascoli, G. A. et al. (2008) ‘Petilla terminology: Nomenclature of features of GABAergic interneurons of the cerebral cortex’, Nature Reviews Neuroscience, 9(7), pp. 557–568. doi: 10.1038/nrn2402.

Babaev, O. et al. (2016) ‘Neuroligin 2 deletion alters inhibitory synapse function and anxiety-associated neuronal activation in the amygdala’, Neuropharmacology, 100, pp. 56–65. doi: 10.1016/j.neuropharm.2015.06.016.

Blundell, J. et al. (2009) ‘Increased anxiety-like behavior in mice lacking the inhibitory synapse cell adhesion molecule neuroligin 2’, *Genes*, Brain and Behavior, 8(1), pp. 114–126. doi: 10.1111/j.1601-183X.2008.00455.x.

Budreck, E. C. and Scheiffele, P. (2007) ‘Neuroligin-3 is a neuronal adhesion protein at GABAergic and glutamatergic synapses’, European Journal of Neuroscience, 26(7), pp. 1738–1748. doi: 10.1111/J.1460-9568.2007.05842.X/FORMAT/PDF.

Cao, F. et al. (2020) ‘Neuroligin 2 regulates absence seizures and behavioral arrests through GABAergic transmission within the thalamocortical circuitry’, Nature Communications. Springer US, 11(1). doi: 10.1038/s41467-020-17560-3.

Chang, E. F. and Merzenich, M. M. (2003) ‘Environmental Noise Retards Auditory Cortical Development’, Science, 300(April), pp. 498–502.

Chao, H.-T. et al. (2010) ‘Dysfunction in GABA signalling mediates autism-like stereotypies and Rett syndrome phenotypes.’, Nature, 468(7321), pp. 263–9. doi: 10.1038/nature09582.

Chen, C.-H. et al. (2017) ‘Neuroligin 2 R215H Mutant Mice Manifest Anxiety, Increased Prepulse Inhibition, and Impaired Spatial Learning and Memory’, Frontiers in Psychiatry, 8, p. 257. doi: 10.3389/fpsyt.2017.00257.

Chih, B., Engelman, H. and Scheiffele, P. (2005) ‘Control of Excitatory and Inhibitory Synapse Formation by Neuroligins’, Science, 307(22), pp. 1324–1328. doi: 10.1126/science.1103773.

Chubykin, A. A. et al. (2007) ‘Activity-Dependent Validation of Excitatory versus Inhibitory Synapses by Neuroligin-1 versus Neuroligin-2’, Neuron, 54, pp. 919–931. doi: 10.1016/j.neuron.2007.05.029.

Gibson, J. R., Huber, K. M. and Südhof, T. C. (2009) ‘Neuroligin-2 Deletion Selectively Decreases Inhibitory Synaptic Transmission Originating from Fast-Spiking but Not from Somatostatin-Positive Interneurons’, Journal of Neuroscience, 29(44), pp. 13883–13897.

Horn, M. E. et al. (2017) ‘Somatostatin and parvalbumin inhibitory synapses onto hippocampal pyramidal neurons are regulated by distinct mechanisms’, PNAS. doi: 10.1073/pnas.1719523115.

Jiang, D.-Y. et al. (2018) ‘GABAergic deficits and schizophrenia-like behaviors in a mouse model carrying patient-derived neuroligin-2 R215H mutation’, Molecular Brain, 11(31). doi: 10.1186/s13041-018-0375-6.

Liang, J. et al. (2015) ‘Conditional neuroligin-2 knockout in adult medial prefrontal cortex links chronic changes in synaptic inhibition to cognitive impairments’, Molecular Psychiatry, 20, pp. 850–859. doi: 10.1038/mp.2015.31.

Messier, J. E. et al. (2018) ‘Targeting light-gated chloride channels to neuronal somatodendritic domain reduces their excitatory effect in the axon’, eLife, 7, pp. 1–21. doi: 10.7554/eLife.38506.

Parente, D. J. et al. (2017) ‘Neuroligin 2 nonsense variant associated with anxiety, autism, intellectual disability, hyperphagia, and obesity’, American Journal of Medical Genetics, Part A, 173(1), pp. 213–216. doi: 10.1002/ajmg.a.37977.

Pfeffer, C. K. et al. (2013) ‘Inhibition of inhibition in visual cortex: the logic of connections between molecularly distinct interneurons’, Nature Neuroscience, 16(8), pp. 1068–1076. doi: 10.1038/nn.3446.

Poulopoulos, A. et al. (2009) ‘Neuroligin 2 Drives Postsynaptic Assembly at Perisomatic Inhibitory Synapses through Gephyrin and Collybistin’, Neuron, 63, pp. 628–642. doi: 10.1016/j.neuron.2009.08.023.

Rubenstein, J. L. R. and Merzenich, M. M. (2003) ‘Model of autism: Increased ratio of excitation/inhibition in key neural systems’, Genes, Brain and Behavior, 2(5), pp. 255–267. doi: 10.1034/j.1601-183X.2003.00037.x.

Rudy, B. et al. (2011) ‘Three groups of interneurons account for nearly 100% of neocortical GABAergic neurons’, Developmental Neurobiology, 71(1), pp. 45–61. doi: 10.1002/dneu.20853.

Schuler, V. et al. (2001) ‘Epilepsy, hyperalgesia, impaired memory, and loss of pre- and postsynaptic GABAB responses in mice lacking GABAB(1)’, Neuron, 31(1), pp. 47–58. doi: 10.1016/S0896-6273(01)00345-2.

Selimbeyoglu, A. et al. (2017) ‘Modulation of prefrontal cortex excitation / inhibition balance rescues social behavior in CNTNAP2-deficient mice’.

Seok, B. S. et al. (2018) ‘The effect of Neuroligin-2 absence on sleep architecture and electroencephalographic activity in mice’, *Molecular Brain*. Molecular Brain, 11(52), pp. 1–11.

Staiger, J. F. et al. (2004) ‘Calbindin-Containing Interneurons Are a Target for VIP-Immunoreactive Synapses in Rat Primary Somatosensory Cortex’, Journal of Comparative Neurology, 468(2), pp. 179–189. doi: 10.1002/cne.10953.

Staiger, J. F. and Petersen, C. C. H. (2021) ‘Neuronal circuits in barrel cortex for whisker sensory perception’, Physiological Reviews, 101(1), pp. 353–415. doi: 10.1152/physrev.00019.2019.

Takács, V. T., Freund, T. F. and Nyiri, G. (2013) ‘Neuroligin 2 Is Expressed in Synapses Established by Cholinergic Cells in the Mouse Brain’, PLoS ONE, 8(9). doi: 10.1371/journal.pone.0072450.

Taniguchi, H. et al. (2011) ‘A Resource of Cre Driver Lines for Genetic Targeting of GABAergic Neurons in Cerebral Cortex’, Neuron, 71, pp. 995–1013. doi: 10.1016/j.neuron.2011.07.026.

Tremblay, R., Lee, S. and Rudy, B. (2016) ‘GABAergic Interneurons in the Neocortex: From Cellular Properties to Circuits’, Neuron, 91, pp. 260–292. doi: 10.1016/j.neuron.2016.06.033.

Varoqueaux, F. et al. (2006) ‘Neuroligins Determine Synapse Maturation and Function’, Neuron. Elsevier Inc, 51, pp. 741–754. doi: 10.1016/j.neuron.2006.09.003.

Varoqueaux, F., Jamain, S. and Brose, N. (2004) ‘Neuroligin 2 is exclusively localized to inhibitory synapses’, European Journal of Cell Biology, 83, pp. 449–456. doi: 10.1078/0171-9335-00410.

Vogt, B. A. (1991) The Role of Layer I in Cortical Function, Normal and Altered States of Function. Cerebral Cortex, vol 9. Edited by A. Peters and E. G. Jones. Springer, Boston, MA. doi: 10.1007/978-1-4615-6622-9_2.

Wilson, H. R. and Cowan, J. D. (1972) ‘Excitatory and Inhibitory Interactions in Localized Populations of Model Neurons’, Biophysical Journal. Elsevier, 12, pp. 1–24. doi: 10.1016/S0006-3495(72)86068-5.

Wöhr, M. et al. (2013) ‘Developmental delays and reduced pup ultrasonic vocalizations but normal sociability in mice lacking the postsynaptic cell adhesion protein neuroligin2’, Behavioural Brain Research, 251, pp. 50–64. doi: 10.1016/j.bbr.2012.07.024.

Xu, Haifeng et al. (2019) ‘A Disinhibitory Microcircuit Mediates Conditioned Social Fear in the Prefrontal Cortex’, Neuron. Elsevier Inc., 102(3), pp. 668–682.e5. doi: 10.1016/j.neuron.2019.02.026.

Xue, M., Atallah, B. V and Scanziani, M. (2014) ‘Equalizing excitation–inhibition ratios across visual cortical neurons’, Nature, 511, pp. 596–600. doi: 10.1038/nature13321.

Yaylaoglu, M. B. et al. (2005) ‘Comprehensive expression atlas of fibroblast growth factors and their receptors generated by a novel robotic in situ hybridization platform’, Developmental Dynamics, 234(2), pp. 371–386. doi: 10.1002/dvdy.20441.

Yizhar, O. et al. (2011) ‘Neocortical excitation/inhibition balance in information processing and social dysfunction’, Nature. Nature Publishing Group, 477(7363), pp. 171–178. doi: 10.1038/nature10360.

Zhang, B. et al. (2015) ‘Neuroligins Sculpt Cerebellar Purkinje-Cell Circuits by Differential Control of Distinct Classes of Synapses’, Neuron, 87, pp. 781–796. doi: 10.1016/j.neuron.2015.07.020.

Zhang, B. and Südhof, T. C. (2016) ‘Neuroligins Are Selectively Essential for NMDAR Signaling in Cerebellar Stellate Interneurons’, Journal of Neuroscience, 36(35), pp. 9070–9083. doi: 10.1523/JNEUROSCI.1356-16.2016.

Zhang, L. I., Bao, S. and Merzenich, M. M. (2002) ‘Disruption of primary auditory cortex by synchronous auditory inputs during a critical period’, Proceedings of the National Academy of Sciences of the United States of America, 99(4), pp. 2309–2314. doi: 10.1073/pnas.261707398.

